# A Scale-Free Gradient of Cognitive Resource Disruptions in Childhood Psychopathology

**DOI:** 10.1101/2021.08.24.457554

**Authors:** Andrew J. Stier, Carlos Cardenas-Iniguez, Omid Kardan, Tyler M. Moore, Francisco A. C. Meyer, Monica D. Rosenberg, Antonia N. Kaczkurkin, Benjamin B. Lahey, Marc G. Berman

## Abstract

The Hurst exponent (***H***) isolated in fractal analyses of neuroimaging time-series is implicated broadly in cognition. The connection between ***H*** and the mathematics of criticality makes it a candidate measure of individual differences in cognitive resource allocation. Relationships between ***H*** and multiple mental disorders have been detected, suggesting that ***H*** is transdiagnostically associated with psychopathology. Here, we demonstrate a gradient of decreased ***H*** with increased general psychopathology and attention-deficit/hyperactivity extracted factor scores during a working memory task which predicts concurrent and future working memory performance in 1,839 children. This gradient defines psychological and functional axes which indicate that psychopathology is associated with an imbalance in resource allocation between fronto-parietal and sensory-motor regions, driven by reduced resource allocation to fonto-parietal regions. This suggests the hypothesis that impaired cognitive function associated with psychopathology follows from a reduced cognitive resource pool and a reduction in resources allocated to the task at hand.

## Introduction

Fractals are found everywhere in the natural world. These patterns are pervasive and include the growth of Romanesco broccoli, the shape of coastlines (*1*), and even the time-series of human neuroimaging data (*2–5*). The defining characteristic of fractals is scale-invariance which refers to the fact that fractals looks the same across levels of magnification. In other words, small pieces of fractals are similar to larger pieces, i.e., fractals display self-similarity. In the context of human neuroimaging data, fractalness is commonly estimated by the Hurst exponent (*6*), *H*, and refers to the degree to which the collected time-series (e.g., from blood oxygen level dependent functional magnetic resonance imaging, BOLD fMRI, or electroencephalography, EEG) are scale-free, or self-similar, in time.

Interestingly, *H* measured from human neuroimaging data is associated with diverse aspects of cognition including learning, task difficulty, and typical adult aging. For example, *H* has been shown to distinguish individuals who benefit from practicing a task vs. those who do not benefit from practice (*7*), to distinguish individuals with and without clinical diagnoses of depression (*8*), to correlate with age, task difficulty, and task novelty (*4*), to track working memory effort beyond working memory capacity (*5*), to track facial-encoding task performance (*2*), and has been found to discriminate highly impulsive from less impulsive persons during performance of an inhibition task (*9*). A convergent lesson from these studies is that across the varied aspects of cognition they explore, they all suggest the heuristic that lower *H* is associated with more cognitive effort.

One hypothesis to explain this convergence of findings is that *H* serves as a real-time index of how close the brain is to a critical state (*10*)^1^, which serves as a proxy for the overall optimality of human neural networks (i.e., how well they maximize energy use for neural activity) (*12*). The concept of critical states is an idea borrowed from the physics of complex systems and statistical mechanics that describes systems at points of transition. For example, water at 374 Celsius and 3,200 psi readily moves between a liquid, a gas, and a solid state. For complex networks, like the brain, critical states provide maximum dynamic range (*13, 14*) and optimized information storage and transfer (*15–17*). Brain states closer to a critical state have values of *H* close to 1 and have smooth looking temporal fluctuations; brain states farther from a critical state have values of *H* closer to 0.5 and have temporal fluctuations that look like random noise (Figure 1) (*4*)^2^. One consequence of these properties of complex systems near critical states is that transitions from critical states into task-relevant states are likely easier and may lead to superior task performance. In support of this idea, *H* has been proposed to quantify how hard it is to transition into a brain state and to quantify the cognitive resources available to make those transitions (*2–5*).

**Figure 1:**
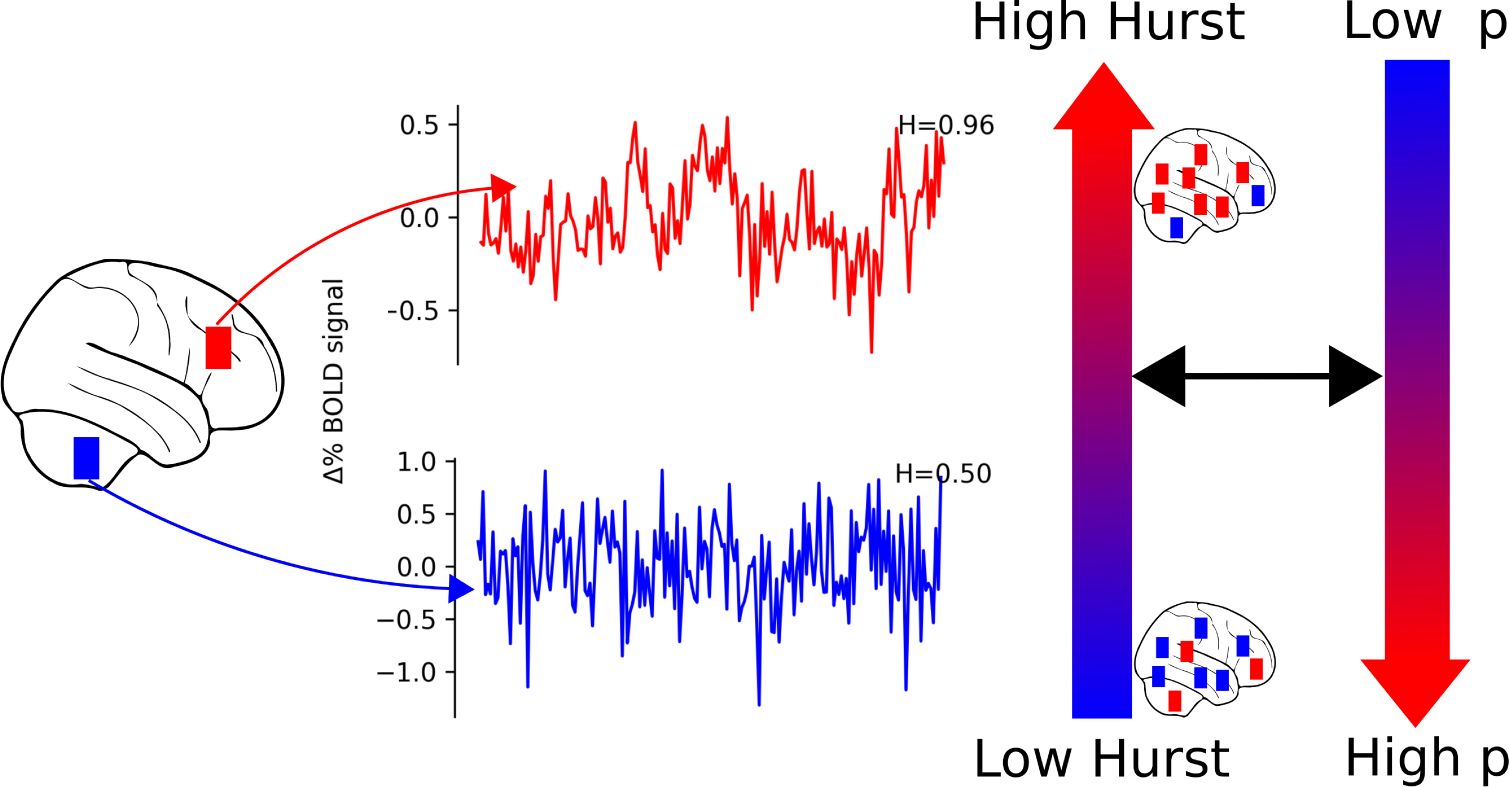
Time-series with *H* close to 1 have smooth looking temporal fluctuations (top). Such time-series can also be described as scale-free or fractal in time. In contrast, time-series with more random temporal fluctuations have *H* closer to 0.5 (bottom). We hypothesized that higher *H* would be associated with lower extracted factor scores for a general factor of psychopathology, p. Though we characterize individuals as having lower *H* or higher *H*, there is still variability in *H* across brain regions.

Since decreased cognitive resources (*18*) and variations in the allocation of cognitive resources among different cognitive systems have been linked to the mental disorder of depression (*19, 20*), individual differences in *H* are of great interest to research on psychopathology. In addition, impulsivity and poor performance on cognitive tasks (which have also been associated with decreased *H* (*2, 9*)) have been linked to risk for essentially all forms of psychopathology (*21*).

Consistent with these findings, *H* calculated from resting state fMRI (*22*) and electroencephalograpic (EEG) data (*23*) have been associated with depression (*8*), cocaine dependence (*24*), attention-deficit/hyperactivity disorder (ADHD) (*25*), schizophrenia (*26*), and autism (*27*). While these studies focused on specific mental disorders, together, these findings suggest that *H* is a non-specific correlate of psychopathology in general. Here, we tested this hypothesis by modeling psychopathology with a bifactor model (*28–31*).

This model asserts that essentially every dimension of psychopathology is positively correlated because they all share causes and psychobiological mechanisms to a considerable degree (*30*). Rather than studying the correlates of every form of psychopathology separately, the bifactor model (*21, 32*) specifies a general factor (also referred to as the a p-factor (*28*)) that reflects transdiagnostic processes contributing to the development of mental disorders in general. This model also specifies orthogonal specific factors that reflect processes that uniquely contribute to subsets of psychiatric disorders. These model specifications match our hypothesis that individual differences in *H* are associated transdiagnostically with psychopathology, thus motivating our use of a bifactor model to characterize psychopathology.

In addition, studies associating *H* with specific aspects psychopathology have not clarified a mechanism which explains these associations. To remedy this, we build on previous literature that has demonstrated 1) decreased *H* under conditions of reduced cognitive resources (e.g., during tasks (*3, 4*), with typical aging in adults (*4*), and in psychopathology (*8, 24–27*)), and, 2) regional variation in the degree to which *H* is reduced during task performance (compared to rest) that may be dependent on the specific demands of the task (*3*). These observations suggest that changes in *H* may originate endogenously (e.g. with age or psychopathology) or exogenously (e.g. via engagement in a task). Importantly, the significance of such changes in *H* vary depending on their source. For example, lower *H* in older individuals has been associated with decreased cognitive performance related to normal aging (*4*). In contrast, in the context of a cognitive task, higher *H* (i.e., a failure to suppress *H*) might indicate a lack of engagement with the task (*2, 5*) and poorer performance.

Consequently, we propose that: 1) if decreased *H* is associated transdiagnostically with psychopathology, it is indicative of reduced global cognitive resources (Figure 1) and 2) relative to an individual’s whole brain pattern of *H*, higher *H* within cognitive brain networks is indicative of relatively fewer cognitive resources allocated toward those cognitive systems, and should also be associated with poorer performance, and vice versa (Figure 2). To help fill these gaps in the literature, we investigated the relationship between *H* during the emotional n-back task (EN-Back, see Methods for details) and psychopathology in a diverse sample of 1,839 children aged 9-10 years old.

**Figure 2:**
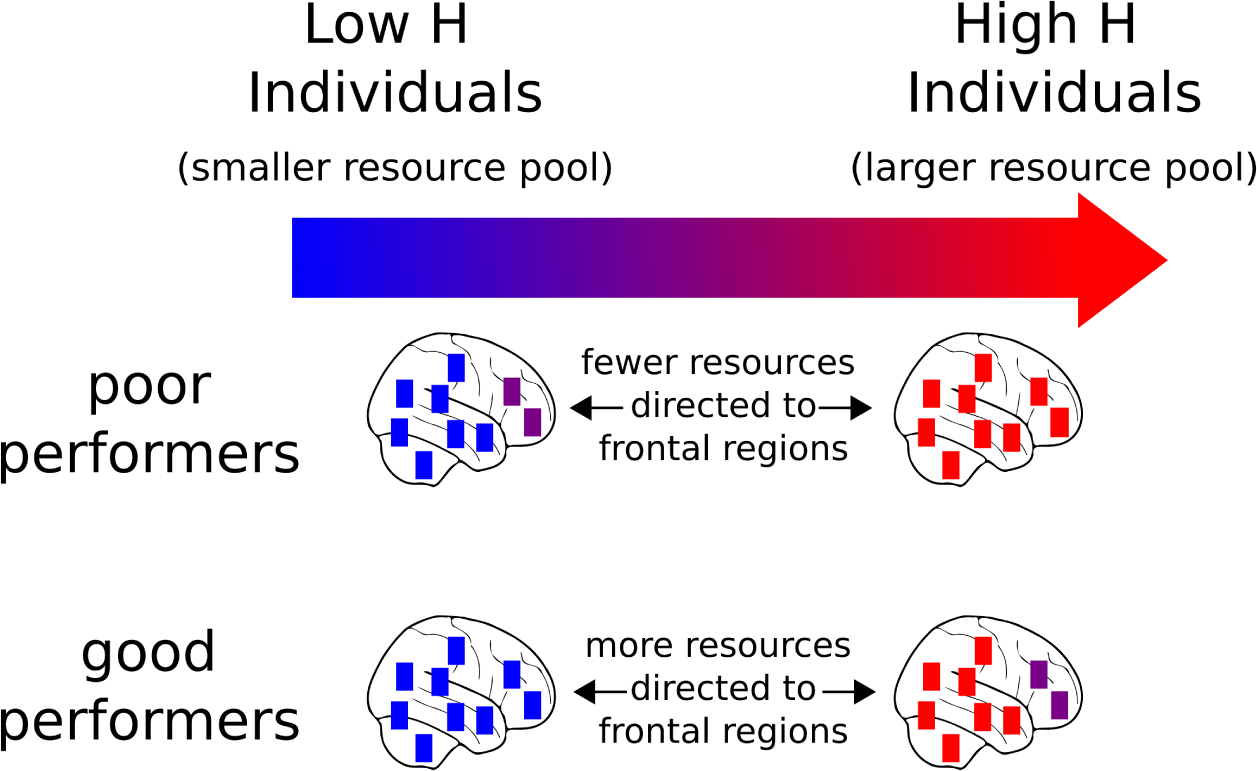
While some individuals differ in overall, whole brain patterns of *H* (see Figure 1), we expect relatively higher *H* in cognitive brain networks to be indicative of fewer resources directed towards the task at hand and poorer task performance, and vice versa. Here we represent this hypothesis by showing the distribution of *H* throughout the brain for individuals with both high *H* and low *H* performing a task which requires recruiting frontal regions. Relative to the whole brain tendency of high or low *H*, we hypothesize that individuals who perform poorly would have higher *H* in frontal regions. This indicates a failure to properly engage and direct cognitive resources to those regions. In contrast, we hypothesize that individuals who perform well, would have lower *H* in frontal regions. This indicates that those individuals engage those regions and direct cognitive resources towards them, resulting in good performance on the task.

## Results

To establish a relationship between *H* and a general factor of childhood psychopathology we used fMRI time-series from the EN-Back task and scores on the parent-rated Child Behavior Checklist (CBCL) from the baseline year of the **Adolescent Brain Cognitive Development***^SM^* **Study** (ABCD) dataset (Release 2.0.1) (*33*). CBCL scores were fit to a bifactor model via confirmatory factor analysis (*21*) (see Methods). Psychopathology was assessed via extracted factors scores from the bifactor model for each subject on the general factor and three specific factors: externalizing, internalizing, and ADHD. After preprocessing, visual quality control, and exclusion for high head motion, the fMRI data from 1,839 children were parcellated into 392 previously defined cortical and subcortical parcels (*34*). *H* for each parcel was then computed (see Methods).

We calculated *H* from each subject’s fMRI BOLD time-series during runs of the EN-Back task, which measures recognition memory in the presence of varying emotional distractors and has been described in detail elsewhere (*33, 35*). Briefly, participants saw a series of images during each block of the EN-Back task and were asked to indicate whether the current image matches the n^th^ previous image. For example, 2-back blocks require remembering images that appeared 2 images before the current image, while 0-back blocks serve as a target detection task in which participants were shown a target image at the beginning of the block and were instructed to respond when the presented image matched the target (*35*). Images consisted of either physical places or faces expressing happy, fearful, or neutral emotions. This task simultaneously probes emotional regulation and working memory, both of which have been implicated transdiagnostically in childhood psychopathology (*36–41*). Consequently, we expected to find lower *H* to be associated with higher extracted bifactor scores (*2, 4*)).

### A Task-Induced Hurst-Psychopathology Gradient

We related *H* to extracted bifactor scores using partial least squares analysis (PLS). PLS is a multivariate technique which extracts maximally co-varying latent variables from two sets of data (*42, 43*) (Figure 3a); in this case *H* in each brain parcel (set 1) with extracted bifactor psychopathology scores (set 2). These latent variables (LVs) consist of loadings on to each of the two sets of data which specify the contribution of data variables (e.g., *H* in occipital cortex) to the LV. Statistically, the LV as a whole is assessed for significance with permutation testing, and the influence of individual variables (e.g., extracted general factor scores or *H* in occipital cortex) is assessed via bootstrap resampling. Importantly, this analysis allows for a direct, multivariate, association between the temporal dynamics of the BOLD signal and psychopathology.

This analysis revealed a single statistically significant latent variable relating *H* with childhood psychopathology (*p* = 0.0007, 10,000 permutations) that captures 49% of the covariance between *H* and extracted bifactor scores. This latent variable has stable positive loadings on the general and ADHD factors and stable negative loadings onto *H* in the brain, where stability was assessed with bootstrap ratios (Figure 3b, see methods). Only brain areas with negative loadings were stable (absolute value bootstrap ratios *>* 3, see Supplementary Table 7). This pattern of psychopathology loadings and brain loadings represents a Hurst-Psychopathology gradient (H-P gradient) associating higher extracted bifactor scores (notably the general factor of psychopathology and ADHD) with lower *H*. In addition, this fits the heuristic supported by past research of lower *H* being associated with more effort, where psychopathology is understood to be a more effortful state (in the colloquial sense).

**Figure 3:**
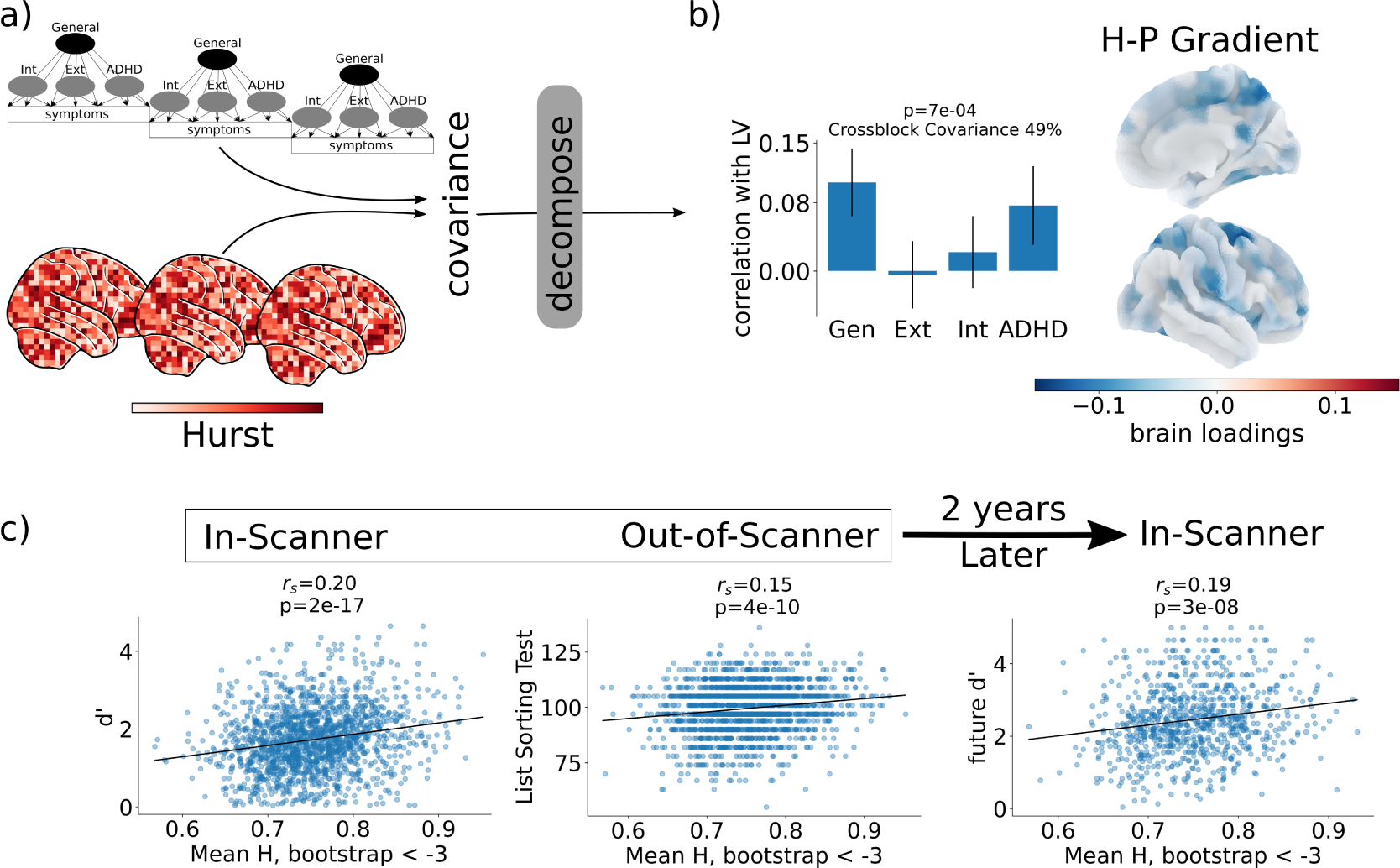
A Task-induced gradient relating scale-free brain activity as measured by *H* to childhood psychopathology. a) We computed the covariance matrix between a subjects’ *H* during the EN-back task and their extracted factor scores as defined in the bifactor model. This covariance matrix was decomposed with partial least squares analysis to find maximally co-varying latent-variables. b) A single latent-variable, which explained 49% of the crossblock covariance was significant. This LV describes a Hurst-Psychopathology gradient in which lower *H* is associated with both higher General and ADHD extracted factor scores. Significance was assessed via 10,000 random permutations. Stability was assessed with bootstrap ratios calculated as the empirical loading divided by the bootstrap variance and is distributed normally under the null (akin to a z-score); 10,000 bootstrap re-samples were used. Results were similar when controlling for family membership (randomly keeping one family member) and when controlling for non-participation (which individuals could complete the task or had minimal head movement) and post-stratification (correcting to nationally representative demographics); see Figure S1. c) Mean *H* for brain regions with absolute value bootstrap ratios *>* 3 are positively correlated with both in-scanner (left) and out-of-scanner (middle) and future-in-scanner (right) working memory performance. This indicates that individuals with lower *H* tend to have higher extracted general and ADHD bifactor scores and also worse working memory performance, and that these deficits persist over time (years) and across tasks (the in scanner and out of scanner memory tasks were different, i.e., EN-back task vs. the List Sorting working memory task).

To increase our certainty that these associations would replicate beyond the characteristics (e.g. socio-demographic factors or scanner manufacturer) of the ABCD Study® sample (*44, 45*), we ran a number of sensitivity tests (see Methods). These tests demonstrated that the H-P gradient was robust to exclusion of multiple family members (randomly retaining only one participant per family), to conditioning on acquisition site and scanner, to the application of post-stratification (to account for differences between the full ABCD sample and nationally representative socio-demographics) and non-participation weights (to account for which children were able to complete the EN-Back task with low head motion), and to the use of CBCL syndrome scales in place of extracted bifactor scores; see Methods, Supplementary Table 1, Supplementary Figures 1 and 2. Consequently, subsequent analyses make use of the un-adjusted H-P gradient.

### Relationship with Task Performance

Next, we asked whether the H-P gradient was associated with working memory performance. In order to do so we averaged *H* over a globally distributed (see Figure 3b) network of brain regions composed of parcels with stable brain loadings (absolute value bootstrap ratios *>* 3). Task performance was assessed in-scanner by sensitivity, d’, and accuracy both during the 2-back blocks of the EN-Back task, and out-of-scanner by un-adjusted, standardized scores on the List Sorting working memory task from the NIH cognitive toolbox (*46*). We note that while 2-back accuracy and d’ are highly correlated (*r_s_* = 0.87, *p* = 0.0), accuracy in this context is strongly influenced by true negatives (correct rejections) while d’ better captures the balance between hits (true positives) and false alarms (false positives). Mean *H* in this distributed stable network was significantly correlated to in-scanner performance (for d’, *r_s_* = 0.20, *p* = 2*e −* 17; for acc, *r_s_* = 0.25, *p* = 2*e −* 25) and out-of-scanner performance (*r_s_* = 0.15, *p* = 4*e −* 10), Figure 3c, indicating that higher *H* is related to better performance both in scanner and out of scanner. Mean *H* in this network was also significantly related to out-of-scanner performance when controlling for in-scanner performance (Supplementary Table 2), suggesting that the H-P gradient is related to working memory ability in a trait-like manner. These results are in line with our hypothesis that decreased *H* associated transdiagnostically with psychopathology is indicative of reduced global cognitive resources (Figure 1).

Additionally, a mediation analysis (see Methods) indicated that mean *H* in this H-P gradient network partially mediates the relationship between extracted bifactor scores and in-scanner working memory performance (Supplementary Table 5). When extracted bifactor scores are treated as the mediator instead of *H* (Supplementary Table), the significant paths remain the same, however, there is no longer a strong relationship between extracted bifactor scores and 2-back accuracy. Thus, while there is no statistical preference for *H* over extracted bifactor scores as a mediator, we can conclude that *H* has a more direct relationship with 2-back working memory performance than psychopathology.

Next, we assessed the relationship between *H* in this H-P gradient brain network and future in-scanner 2-back d’ and accuracy during the 2-year follow-up session. Specifically, we correlated the same baseline H-P gradient measure for each participant with in-scanner 2-back accuracy and d’ during the 2-year follow-up session for participants that had data for both sessions (N=888, approximately half of the full sample had 2-year follow-up data available in ABCD Release 3.0). Since the sample of individuals who had available data for ABCD Release 3.0 is non-random (see Supplementary Table 3), we additionally computed weighted correlations to correct for non-participation (see Methods). Only future d’ was significantly correlated with the H-P gradient measure after correction for non-participation (*r_s_* = 0.19, *p* = 3*e −* 8; 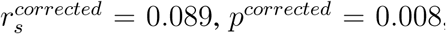, Figure 3c). Before correction 2-back accuracy was similarly correlated with the H-P gradient measure (*r_s_* = 0.20, *p* = 8*e −* 11), but this was not significant after correction 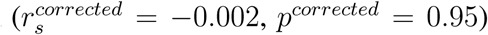. In addition, though baseline in-scanner 2-back d’ and 2-year follow-up in-scanner 2-back d’ are significantly correlated (*r_s_* = 0.51, *p* = 4*e −* 57), mean *H* in the H-P gradient network was significantly related to future d’ when controlling for baseline d’ (Supplementary Table 4).

In summary, these results indicate that the Hurst-Psychopathology gradient is associated with trait-like deficits in working memory that persist over long time periods (i.e., years) and are seen in a variety of working memory tasks.

### The H-P Gradient Defines a Functional Activation Axis

We next sought to better understand the meaning of the H-P gradient and to understand its context in the larger body of literature investigating functional activations, task performance, and psychopathology. To do so we determined whether there were patterns of functional activation within large scale cognitive systems associated with the H-P gradient.

We examined the relationship between individual subjects’ H-P associations and their own simultaneous functional activations (i.e., BOLD signal contrast during the same run). Specifically, we asked how each subject’s pattern of activations during the EN-Back task was related to the degree to which they exemplify the H-P gradient. To do so we generated a gradient score for each subject by correlating their Hurst map with the H-P gradient map. Higher gradient scores are statistically indicative of higher levels of psychopathology, lower *H*, and poorer task/cognitive performance. Next, activations were defined as the contrast between 2-back and 0-back blocks in 148 cortical regions (see Figure 4a, Methods (*33*), we additionally examined the contrast between emotional and neutral faces, positive and neutral faces, and negative and neutral faces, but there were no significant associations for these contrasts with the H-P gradient after multiple comparison correction; Supplementary Figures 3-5). Finally, we correlated the activation values and subject gradient scores in each of the 148 cortical regions. A positive correlation implies that increased activation (2-back vs 0-back contrast) is associated with higher extracted bifactor scores, and a negative correlation implies that decreased activation is associated with higher extracted bifactor scores.

**Figure 4:**
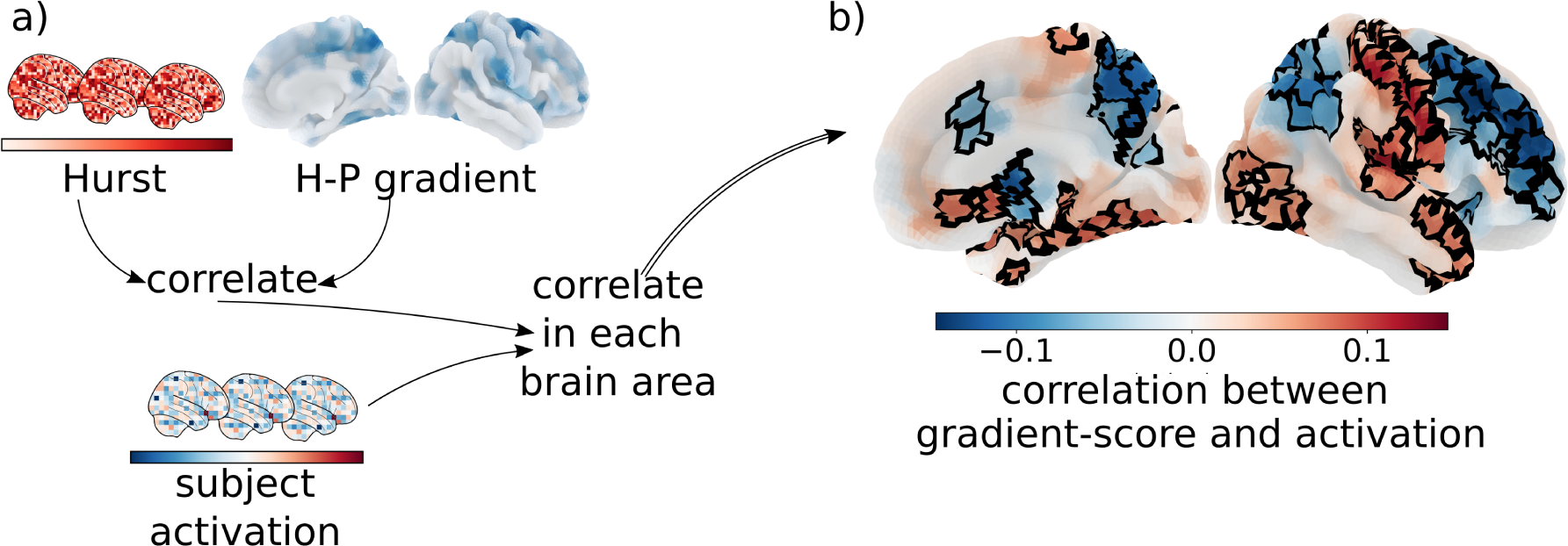
The Hurst-Psychopathology gradient defines functional-activation axis associated with task performance and psychopathology. a) To understand the relationship between the H-P gradient and functional activation we first computed a gradient score for each subject by correlating their spatial patterns of *H* to the H-P gradient. Next, we correlated these gradient scores across subjects to functional activation in each brain parcel. b) Higher general factor of psychopathology and ADHD extracted factor scores and lower H are associated with decreased fronto-parietal activation and increased occipital, medial-temporal, and sensory-motor activation on 2-back vs. 0-back blocks. Areas where gradient scores are significantly correlated with functional activations after multiple comparison correction are outlined in black.

**Figure 5:**
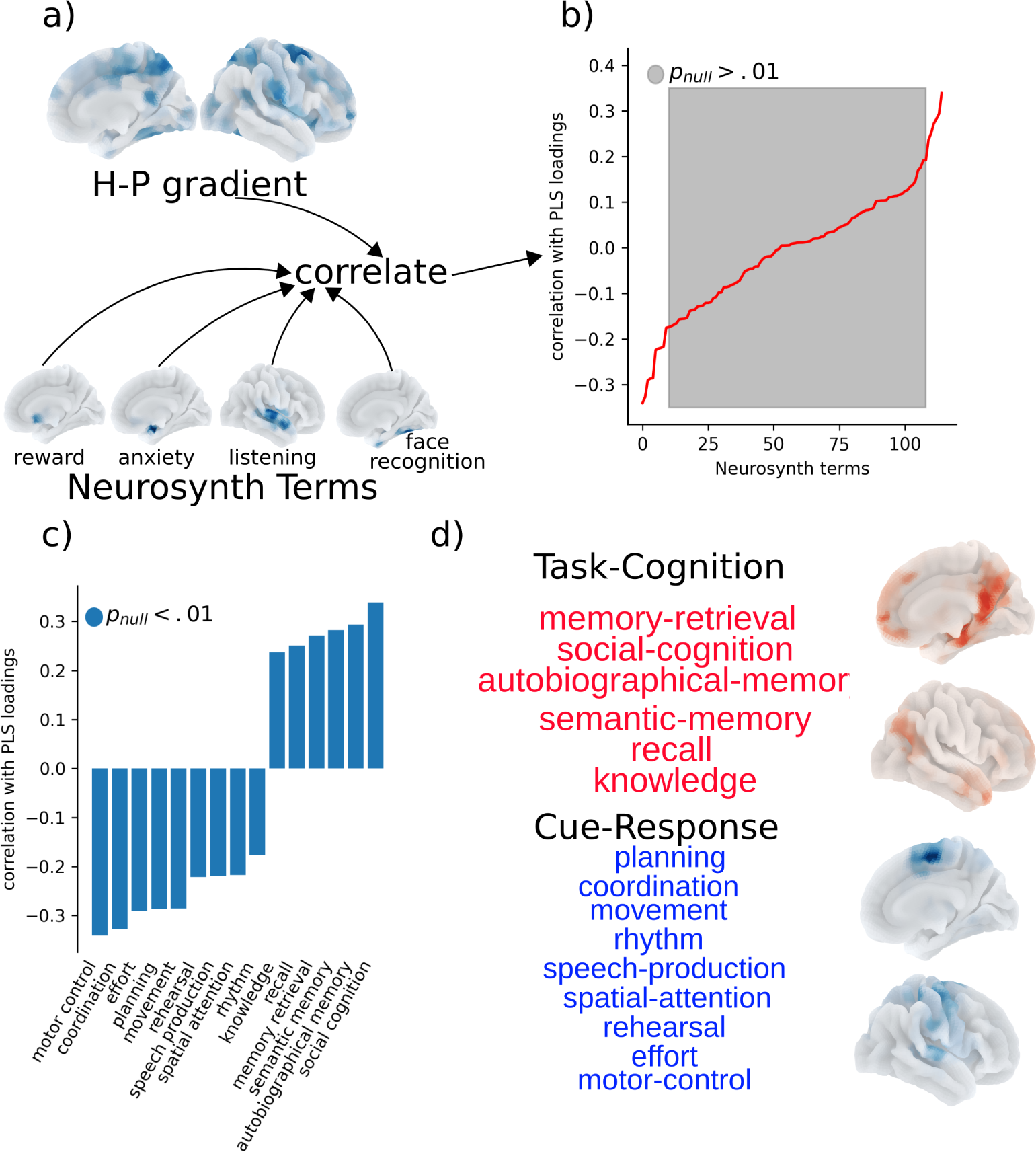
The Hurst-Psychopathology gradient defines a psychological axis associated with task performance and psychopathology. a) We used Neurosynth term association maps showing the probability of activation with multiple psychological terms. The term set was restricted to the intersection of terms used in (*48, 50*) and terms that had available association maps on the Neurosynth website, yielding 116 terms. Each map was correlated with the H-P gradient to identify which terms had spatial patterns of activation most similar to the gradient. b) Grey indicates non-significance based on 1,000 parametric spatial permutation tests (Benjamini-Hochberg correction, *α* = .01). Terms are ranked by magnitudes of correlations. c) Terms that are positively correlated with the H-P gradient are the positive term set and terms that are negatively correlated with the H-P gradient are the negative term set. d) The positively correlated terms include task-relevant cognitive processes. The negatively correlated terms included processes involved in planning and executing responses to task cues. Surface maps of these two axes were created by taking the maximum value across term maps included in each axis.

The resulting map (Figure 4b) indicates that higher gradient scores, (i.e., higher general and ADHD extracted bifactor scores and lower *H*), and poorer working memory performance are associated with decreased fronto-parietal activation and increased sensory-motor activation in the same task. This is in line with previous work in this sample (*35*) which found that lower performance was associated with decreased functional activation (2-back vs. 0-back) in fronto-parietal areas commonly associated with cognitive processes related to this working memory and emotional regulation task. This convergence of findings suggests overlap among the mechanisms driving poorer task performance generally, and the mechanisms driving psychopathology specifically. In other words, this pattern of functional activations– which represents a fronto-parietal/sensory-motor axis of resource use – is not specific to psychopathology. However, its association with psychopathology and *H* here suggests that the general factor of psychopathology and the ADHD specific factor are associated with a relative redistribution of cognitive resources away from task-relevant brain networks and into sensory motor processing.

### The H-P Gradient Defines a Psychological Axis

Next, we sought to understand the significance of spatial variation in the strength of the H-P gradient (i.e., the relationship between Hurst exponents and psychopathology) across different brain regions. For example, this relationship is stronger in some parts of insular cortex and in middle frontal gyrus (two areas associated with emotional regulation and working memory, respectively) but weaker in some parts of occipital cortex and anterior frontal cortex (Supplementary Table 7). Consequently, we sought to better understand the psychological and cognitive significance of these spatial variations within the H-P gradient. In other words, we sought to characterize the data-derived H-P gradient in reference to the broader neuroimaging literature independent of the operationalizations of the present study to increase interpretability.

For this analysis, we first examined similarities between the H-P gradient and probabilistic meta-analysis maps from Neurosynth (*47*) which describe how frequently journal articles contain specific terms alongside voxel coordinates related to functional activation. We expected this analysis to reveal a psychological axis defined by the H-P gradient that contrasts psychological terms which are similar and dis-similar to the H-P gradient.

We used a previously defined subset of 125 terms (*48*) which included, for example, ”cognitive control”, ”language comprehension”, ”memory”, ”psychosis”, and ”social cognition”. This term set was restricted to the overlap between Neurosynth terms and Cognitive Atlas (*49*) terms and can thus be thought of as belonging to a proposed classification of psychological concepts and tasks. Each term’s associated activation map was correlated with the H-P gradient and significance was assessed via a spatial null model (see Methods, Figure 5a). Importantly, these Neurosynth maps only indicate which regions are commonly reported alongside a given term and do not address the sign of the association (i.e., whether the psychological term is associated with functional activation or deactivation).

After correction for multiple comparisons, 15 terms remained (Figure 5b&c). The set of terms positively correlated to the H-P gradient define a Task-Cognition category which includes terms related to the cognitive processes engaged by the EN-Back task (*35*): emotional-regulation and working-memory (Figure 5c&d, top). The set of terms negatively correlated to the H-P gradient define a Cue-Response category which includes terms related to processes involved in planning and executing responses to the task cues (5c&d, bottom). These results describe a Task-Cognition/Cue-Response axis of cognitive function which is relevant to childhood psychopathology.

Finally, we sought to confirm our hypothesis that this Task-Cognition/Cue-Response axis indicates that while individuals with higher general factor and specific ADHD extracted bifactor scores tend to have lower *H* across the whole brain, they also tend to have higher *H* in Task-Cognition areas and lower *H* in Cue-Response areas, relative to their own whole brain pattern of *H*. To do so, we took the union across term maps in the Task-Cognition and Cue-Response sets by choosing the maximum z-score in each brain region. Next, we kept regions above a z-score threshold and calculated the difference between the mean *H* for Task-Cognition and Cue-Response. Finally, we correlated this difference across subjects to gradient scores (i.e., the degree to which each subject exemplifies the H-P gradient during the EN-back task). This resulted in a significant positive correlation across all choices of Z-score threshold (Supplementary Figure 6). Since higher gradient scores are statistically indicative of higher psychopathology and poorer working memory performance, this result indicates that psychopathology is associated with increases in *H* in Task-Cognition areas and decreases in *H* in Cue-Response areas, relative to individuals’ own whole brain *H*.

**Figure 6:**
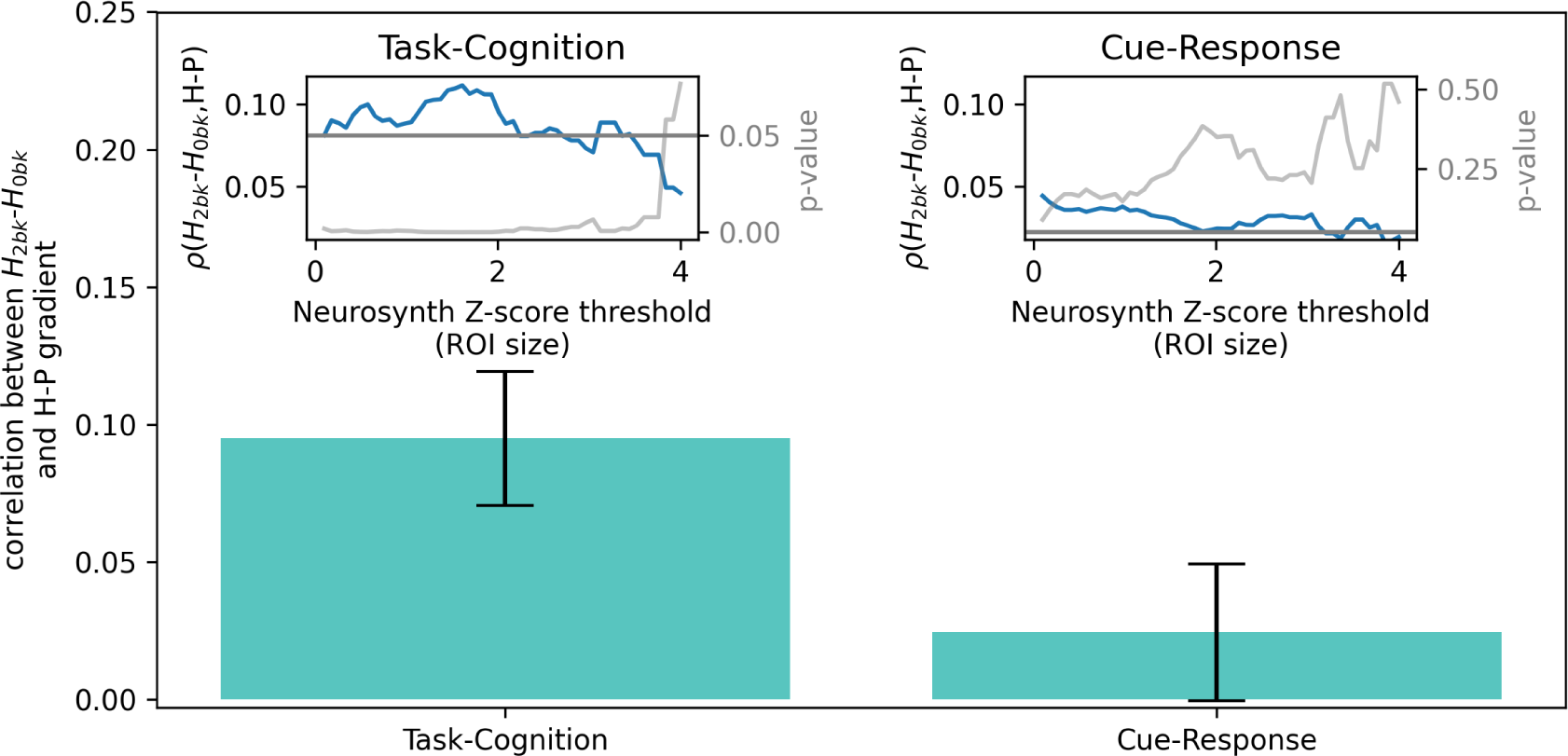
Correlations (blue) were computed across all choices of Z-score threshold used to define the task-cognition and cue-response ROIs from Neurosynth meta-analysis probabilistic activation maps. Significance of the correlations were assessed at all choices of threshold (gray). The p=0.05 level is show as the gray horizontal line. We found evidence of a correlation between higher *H* and higher H-P gradient scores only for task-cognition areas. Insets show correlations between block level 2-vs. 0-back *H* contrast and H-P gradient scores for all choices of Z-score threshold. The bar plot shows the correlation for the choice of Z=2. Error bars represent standard deviations from 1000 bootstrap resamples.

Further, this relative difference in *H* between Task-Cognition and Cue-Response areas tends to be exacerbated in individuals with higher general and ADHD extracted bifactor scores, as demonstrated by positive correlations in Supplementary Figure 6. This parallels the functional fronto-parietal/sensory-motor axis which indicated that psychopathology is associated with reduced functional activation in fronto-parietal areas and increased functional activation in sensory-motor areas.

More specifically, the fronto-parietal/sensory-motor functional axis defined by the H-P gradient identifies two networks of brain areas in which reduced/increased functional activation is associated with psychopathology. These areas (Figure 4) broadly fit into two groups which are associated with task-related and sensory-motor processes, respectively. Similarly, the Task-Cognition/Cue-Response psychological axis defined by the H-P gradient identifies two sets of psychological terms (Figure 5) which are associated with similar sets of brain areas as are identified by the functional axis.

Taken together, these two parallel psychological and functional axes suggest that psychopathology is associated with a reduction in resource allocation to Task-Cognition areas, relative to Cue-Response areas. In other words, individuals with higher levels of psychopathology tend engage relatively fewer fronto-parietal resources relevant for the emotional regulation and working memory demands of the EN-Back task and also tend to engage relatively more sensory-motor resources relevant for the rote sensory and motor demands of the task.

### Associations of the H-P gradient with block level Hurst exponents

The previous analyses implicitly contrasted Task-Cognition/fronto-parietal (TC/fp) and Cue-Response/sensory-motor (CR/sm) areas. As a result, those analyses established that psychopathology is associated with a redistribution of resources away from TC/fp regions, relative to CR/sm regions. However, those analyses did not establish whether TC/fp regions or CR/sm regions drive this redistribution of resources associated with psychopathology. Thus, we sought to determine whether a net decrease in resources allocated towards TC/fp regions, a net increase in resources allocated towards CR/sm regions, or a combination of both, was responsible for this effect. In other words, we sought to better understand which brain regions drive the relative redistribution of resources away from TC/fp regions seen in the previous analyses.

To this end, we calculated *H* for 2-back and 0-back blocks of the EN-Back task separately and then averaged *H* across all blocks of the same type for each subject (see Methods, (*51*)). We averaged *H* within the TC/fp and CR/sm regions at various Z-score thresholds on the Neurosynth maps and subtracted 0-back average TC/fp and CR/sm *H* from 2-back average TC/fp and CR/sm *H*, respectively. This created a contrast between 2-back and 0-back blocks within TC/fp and CR/sm areas which would indicate the degree to which *H* was suppressed (*4*) during 2-back blocks, relative to 0-back blocks. Finally, we correlated this contrast with 2-back accuracy (acc), sensitivity (d’), and participant H-P gradient scores.

As expected, irrespective of psychopathology, suppression of *H* during 2-back in TC/fp and CR/sm areas (i.e., lower Hurst during 2-back compared to 0-back) was significantly associated with superior performance as measured by d’ (Supplementary Figure 8; for CR/sm regions p *<* .05 for Neurosynth Z-score threshold, Z, *<* 3.44; for TC/fp regions p *<* .05 for Z *<* 1.85). This effect was driven by hit rate (the percent of responses that are correct) rather than the false alarm rate (Supplementary Figure 9). However, only decreased *H* during 2-back in the TC/fp regions were significantly associated with 2-back accuracy (Supplementary Figure 10, for TC/fp regions p *<* .05 for Z *<* 1.61). These observations further support our hypothesis that relative to individuals’ own whole brain patterns of *H*, higher *H* within cognitive brain networks are indicative of fewer cognitive resources allocated toward those cognitive systems, and vice versa.

In contrast, psychopathology, assessed via higher H-P gradient scores, was significantly associated with less suppression of *H* during 2-back (i.e., less decrease in *H* during 2-back relative to 0-back) in TC/fp areas, but not in CR/sm areas (Figure 6; for TC/fp regions p *<* .05 for Z *<* 3.76). This indicates that psychopathology is related to less engagement of TC/fp regions. Thus, while better task performance generally was associated with suppression of *H* in both TC/fp and CR/sm regions, we only found evidence that psychopathology and associated poorer task performance was associated with higher *H* in TC/fp regions (i.e., less suppression of *H*, see Figure 2). This combination of results suggests that decreased resource allocation towards Task-Cognition/fronto-parietal (TC/fp) regions alone drives the relative redistribution of resources away from TC/fp regions associated with psychopathology.

## Discussion

We found a multivariate pattern (gradient) associating decreased *H* with increased extracted bifactor scores for the general factor of psychopathology and specific ADHD factor, thereby demonstrating for the first time, a general, non-specific association between *H* and psychopathology. Previous research has demonstrated associations between the *H* and specific mental disorders primarily in small populations (*8, 24–27,52, 53*), but did not clarify whether the Hurst exponent was also associated transdiagnostically with psychopathology in large diverse samples.

While the overall pattern observed was that of decreased *H* in individuals with higher extracted general factor and ADHD bifactor scores, there was spatial variation in the strength of this pattern across the brain. To better understand the cognitive and psychological relevance of these spatial variations, we compared this gradient to simultaneous functional activations in the same individuals (*33*) and to meta-analysis-derived activation maps associated with psychological and cognitive terms (*47, 48, 50*).

These analyses revealed that the gradient associating *H* and psychopathology defines a Cue-Response/Task-Cognition psychological axis and a parallel sensory-motor/fronto-parietal activation axis. These axes suggest that individuals with higher extracted bifactor scores tend to allocate fewer resources to Task-Cognition/fronto-parietal areas, measured via reduced functional activation (i.e., BOLD contrast) and relatively higher *H*. In addition, block level analyses suggested that this effect was driven solely by a reduction in resources (measured via higher Hurst exponents) to Task-Cognition/fronto-parietal areas. Together, these results suggest a mechanistic account of reduced task performance associated with psychopathology. Specifically, psychopathology, in general, is associated with an overall reduction in cognitive resources, fewer resources directed towards task-specific cognition, and subsequent poorer task performance. Interestingly, these results also suggest that Cue-Response/sensory-motor areas may have priority when it comes to resource allocation.

In addition, the results presented here are in line with the recent evidence that *H* is a quantitative measure of available cognitive resources and/or mental effort (*2–5*). Under this proposal, deviations from a theoretical state of “perfect” rest, which is assumed to be organized near a critical state and to have a Hurst exponent of 1 (*54, 55*), are indicative of decreased cognitive resources and/or increased mental effort. As the demand for additional cognitive resources increases due to task-demands, fatigue, or psychopathology, the brain moves further away from the critical state, reducing the available supply of flexible information processing resources in exchange for more context relevant processing (*10*).

Previous research has suggested that lower *H* is characteristic of harder tasks (*4*) and more difficult versions of the same task (*5*), but has not clarified how Hurst exponents are related to perceived mental effort when exogenous task demands remain constant. If the results presented here were indicative of transient increases in mental effort, we would not expect generalization to different tasks or across time (i.e., to the same task being performed two years after brain activity was measured). However, we found that the Hurst-Psychopathology gradient is predictive of out-of-scanner (a different task) and future (the same task two years later) working memory performance even when controlling for in-scanner and baseline performance, respectively. Thus, the association between the described Hurst-Psychopathology gradient and working memory performance is relatively stable across time and across at least two working memory tasks as scale-free neural signature of psychopathology. However, these results do not, on the whole, clarify whether deficits in cognitive resources precede or follow psychopathology or clarify the time periods over which decreases in available cognitive resources persist. Future work, possibly with the forthcoming longitudinal waves of the ABCD study dataset, might better be able to answer both questions.

Overall, these findings support the hypothesis that *H* provides a quantitative description of suboptimal brain states that are non-specifically associated with all forms of psychopathology. While these findings specifically identified disruptions in resource allocation among cognitive networks relevant to the task at hand, they also suggest a specific, unifying hypothesis. In particular, these findings suggest that global decreases in *H* can quantify reductions in available cognitive resources, while more local, relative fluctuations in *H* quantify the allocation of available cognitive resources to different cognitive systems. This may provide the basis for a mechanistic account of cognitive performance and the consequences of individual differences in cognitive resources.

The analysis of temporal fractals found in human neuroimaging data promises a systematic framework for understanding human cognition. Quantification of these temporal fractals via the Hurst exponent provides an ingress to the well-developed literature of models and mathematics, rooted in criticality, which may help build concrete, mechanistic models of cognition and allow for systematic characterizations of brain states. The need for such models (*56, 57*) and mechanistic accounts of cognition has been an increasingly important goal in the cognitive sciences. We see the theoretical framework of criticality, and measurements of *H*, as adjacent to other contemporary frameworks that together may provide a comprehensive framework for understanding cognition. As such, explorations of the associations between *H* and cognition may help to elucidate what optimal brain states represent and how deviations from those brain states arise due to psychopathology.

## Online Methods

### Adolescent Brain and Cognitive Development Data

#### Data & Preprocessing

We performed functional MRI preprocessing on ABCD Study baseline year emotional N-Back data, which included 10,240 participants. Participants who were scanned on Phillips brand scanners were excluded because of a known error in the phase encoding direction while converting from DICOM to NIFTI format. We downloaded minimally processed structural and function MRI scans from the ABCD data portal (https://nda.nih.gov/abcd). Minimal preprocessing included motion correction, B0 distortion correction, gradient warping correction and resampling to an isotropic space (*33*). Minimally processed data were preprocessed with a custom version of FMRIPREP (*58*), a Nipype (*59*) based tool. Each participant’s structural T1w scan was first defaced with pydeface (*60*). Each T1w (T1-weighted) volume was then corrected for INU (intensity non-uniformity) using N4BiasFieldCorrection v2.1.0 (*61*) and had been previously skull-stripped. Spatial normalization to the MNI152 non-linear 6th generation template, the standard MNI template included with FSL, was performed through nonlinear registration with the antsRegistration tool of ANTs v2.1.0 (*62*), using brain-extracted versions of both T1w volume and template. Brain tissue segmentation of cerebrospinal fluid (CSF), white-matter (WM) and gray-matter (GM) was performed on the brain-extracted T1w using fast (FSL v5.0.9 (*63*)). Functional data co-registered to the corresponding T1w anatomical image using boundary-based registration (*64*) with six degrees of freedom, using flirt (FSL). Motion correcting transformations (based on motion parameters obtained from the minimally processed data), BOLD-to-T1w transformation and T1w-to-template (MNI) warp were concatenated and applied in a single step using antsApplyTransforms (ANTs v2.1.0) using Lanczos interpolation. Physiological noise regressors were extracted and applied from tissue masks, and frame-wise displacement (*65*) was calculated for each functional run using the implementation of Nipype. For more details of the pipeline see https://fmriprep.readthedocs.io.

Following (*66, 67*) we performed a 36 parameter confound regression that included: the time courses of mean CSF signal, mean global signal, mean WM signal, the 6 standard affine motion parameters (x, y, z, pitch, roll and yaw), their squares, their derivatives, and the squared derivatives of these signals. We also simultaneously regressed out linear and quadratic trends to remove drift related signals. This was followed by the application of a bandpass filter with a highpass cutoff of .008 Hz and a lowpass cutoff of .12 Hz via the 3dBandpass command in AFNI (*68*). The cleaned volumetric BOLD images were spatially averaged into the 392 parcel Craddock atlas (*34*). Finally, for Siemens scanners, the first eight volumes were removed because they were used as the multiband reference. For GE scanners running DV25 software, five volumes were removed because the first 12 volumes were used as the multiband reference and then combined into a single volume and saved as the initial TR (leaving a total of five frames to be discarded). For GE scanners running DV26 software, 16 volumes were removed (*35*). Runs included 362 whole-brain volumes after these discarded acquisitions.

Finally, all structural and functional scans were visually inspected to screen for scanner abnormalities, and to assess the accuracy of the registration and tissue segmentation processes. Only subjects with passing structural scans and at least one passing functional scan were included for further analyses.

### Cognitive Task Procedures

The emotional n-back task engages processes related to memory and emotion regulation (*35*). Each session consists of two approximately 5-minute fMRI runs in which participants complete four 0-back (low working memory load) and four 2-back (high working memory load) task blocks. Each task block consists of four types of stimuli: happy, fearful, and neutral face photographs, and place photographs (*35*). Performance was quantified as percent accuracy and sensitivity, d’, on 2-back blocks (*35, 51*). Sensitivity, d’, was computed as

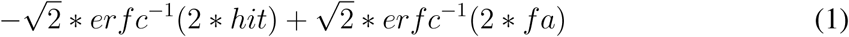

where *erfc* is the complementary error function, *hit* is the hit rate in 2-back blocks, and *fa* is the false alarm rate in 2-back blocks.

### Exclusion of Data

After preprocessing, all fMRI BOLD time courses were spatially averaged within 392 previously defined functional regions (*34*). Individual runs with greater than .2mm mean and 2mm max framewise displacement were excluded, which when combined with visual quality inspections resulted in the retention of 1,839 subjects. For each subject if more than one run was retained, the parcellated time series were averaged over the two runs.

### Estimation of Hurst Exponents

We measured *H* of the mean BOLD time series of each of the 392 previously defined regions using detrended fluctuation analysis (DFA). This is a computationally efficient estimator of the Hurst exponent that is a more robust alternative to power-spectral-density-based methods and has been shown to exhibit convergence with more sophisticated estimators of *H* in fMRI data (*4*). Briefly, DFA involves transforming a detrended time-series, *x*(*t*), into an unbounded random walk, 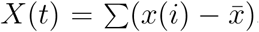, and then calculating local linear fits to this random walk, *Y* (*t*), for various window sizes *n*. The root mean square fluctuations from the local linear trend is then calculated for each window size as 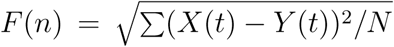, where *N* is the number of windows of size *n*. Finally, *H* is calculated as the slope of a linear fit of *log*(*n*) vs. *log*(*F* (*n*)) (*6, 69*).

For ”mono-fractal” processes, a single exponent *H* fully describes the scaling relationship between local fluctuations and window size. However, in real data, *H* may change as a function of the window size due to the presence of different scaling regimes, or trends in the data that are not controlled by the detrending step of DFA. Thus, it is critical to examine the goodness of fit for the DFA regression line to determine whether such confounds are present (*69*). We examined the goodness of fit using the coefficient of determination, *R*^2^ (Supplementary Figure 7).

We chose to use window sizes, *n*, which exclude possible low-frequency confounds below 0.01 Hz and high frequency confounds above 0.1 Hz. We sampled the number of windows, *N*, approximately uniformly given this frequency constraint. For each number of windows, *N*, we chose the maximum window size *n* such that *n · N* was less than or equal to the number of timepoints in the data.

### Psychopathology measures

#### CBCL Scales

CBCL scales were retrieved from ABCD Release2.0.1 tabulated data available on the NIMH Data Archive (https://nda.nih.gov/abcd). Calculated t-scores for each scale (*70*) were retrieved from the *abcd cbcls01.txt* file. Definitions of the variable names in this file are available in the *abcd cbcls01 definitions.csv* file.

#### Bifactor Scores

The bifactor model was fit using confirmatory factor analyses as in (*21*) and is described there in detail. Briefly, exploratory analyses were conducted on half of the baseline sample to determine which of the 119 CBCL items were most strongly associated with psychopathology. From these reduced set, we extracted four interpretable factors and all CBCL items with a loading *≥* 0.40 on at least one factor were retained. Next, the a confirmatory bifactor model in the second half of the sample was specified based on these results. As required for bifactor models, each retained CBCL item loaded on the general factor and only one specific factor. All other loadings were fixed to zero, and all factors were specified to be orthogonal. Factor models were fit in Mplus 8.3 using the mean- and variance-adjusted weighted least squares (WSLMV) estimator. All factor models accounted for the stratification of the sample in data collection sites, used post-stratification weights, and accounted for clustering within families. Analyses made use of factor score estimates that were extracted from the confirmatory model (*71*).

### Partial Least Squares

Partial least squares (PLS) analysis was used to find a latent variable that represents a Hurst-Psychopathology gradient. PLS is a multivariate data analysis technique that decomposes the covariance matrix between two mean-centered datasets. Here these datasets were the 1,839 by 392, subject by parcel matrix of *H* **X** and the 1,839 by 4, subject by factor matrix of extracted bifactor scores **Y**. Since these matrices are mean-centered, the covariance matrix **X***^t^***Y** can be decomposed via singular value decomposition so that

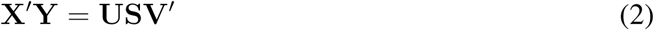

where **U** is the 392 by 4 matrix of left singular vectors (brain loadings), **V** is the 4 by 4 matrix of right singular vectors (bifactor loadings), and **S** is the 4 by 4 diagonal matrix of singular values (*43*). The *i^th^* column of **U** and **V** represent the loadings of the *i^th^* latent variable. The ratio of the squared *i^th^* singular value to the sum of all squared singular values gives the crossblock covariance explained by the *i^th^* latent variable and is used as a measure of effect size.

Statistical significance of the PLS models was assessed by 10,000 permutations of the rows of the Hurst matrix **X** and comparing the observed crossblock covariance to permuted cross-block covariances. Stability of the left and right singular vectors (brain and bifactor loadings, respectively) is assessed by 10,000 bootstrap resamplings of both data matrices, **X** and **Y**.

Bootstrap ratios are calculated as the empirical loading divided by the bootstrap variance and are distributed normally under the null (akin to a z-score).

### Non-Participation, Post-Stratification Weights, and adjustment for Family Membership

Sensitivity analyses were conducted using post-stratification and non-participation weights in an attempt to calibrate ABCD Study sample distributions to nationally representative distributions as measured in the American Community Survey (ACS), and to correct for any biases due to not being included in the analysis relative to the demographic characteristics of the overall ABCD Study sample, respectively. Procedures used to calculate the post-stratification scores (variable *abcd_acs_raked_propensity* in file *acspsw03.txt* of Curated Release 2.0.1) are described in detail elsewhere (*44, 45*). Briefly, a multiple logistic regression model was fit using concatenated ACS and ABCD data to predict study membership using participant variables age, sex, race/ethnicity, family income, family type, household size, parents’ work force status and Census Region. Weights were then raked to exact ACS population counts for age, sex, and race/ethnicity categories. Non-participation weights were derived using an elastic net regularized binary logistic regression model using glmnet in R previously used in child psychopathology studies (*72*). A binary variable indicating inclusion/exclusion in the analysis sample was the dependent variable, while age (in months), sex (male as reference category), race/ethnicity (non-Hispanic white as reference category), household size, years of maternal education, and square-root-transformed mean CBCL score were the independent variables. This elastic net model in glmnet was selected to derive the non-participation weights as it produces estimates with lower predictive errors than the full model, while accounting for redundant and highly correlated potential predictors (*73, 74*). The logistic regression model picked the optimal tuning parameter lambda with the least cross-validation deviance in model selection. Having selected the optimal model, probabilities 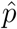 of response, conditional on being sampled, were calculated using the following equation:

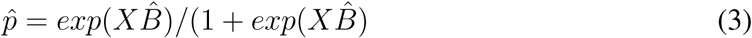

where *X* is the model matrix and 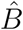 is the vector of estimated parameters from the best model after cross-validation. Non-participation weights are thus the inverses of the probabilities 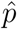. In order to compute corrected correlations between *H* in the H-P gradient and future working memory performance, non-participation weights capturing which individuals had available 2-year follow-up data at the time of this study were also calculated. For the corrected correlations, post-stratification weights, non-participation weights from the full ABCD study sample to this sample of 1,839 children, and non-participation weights from the baseline study sample to the future Release3.0 sample (N=888) were multiplied together.

The non-participation and post-stratification weights were multiplied prior to use in the PLS model. Next the weighted mean, weighted variance, and weighted covariance were computed as (*75*)

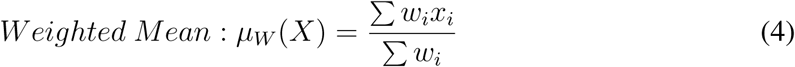

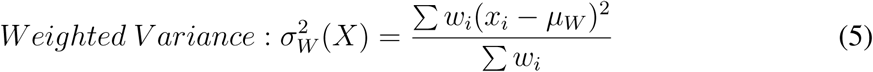

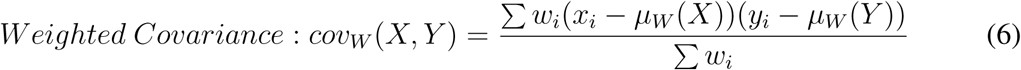

with the resulting weighted covariance matrix then decomposed with singular value decomposition. The corrected correlation between *H* in the H-P gradient and future working memory performance was calculated as:

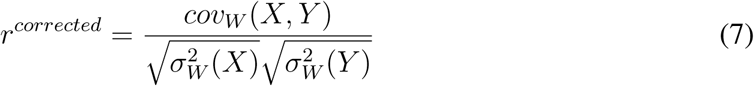

Statistical significance was assessed by computing the weighted mean and variance for the permuted brain matrix and non-permuted bifactor matrix. Next the weighted covariance matrix was calculated as

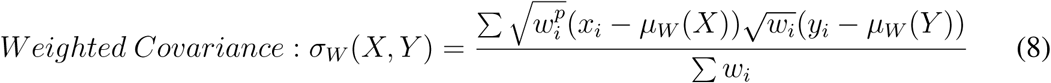

where 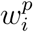 is the weight obtained by permuting the weights alongside the rows of the brain matrix **X.** Finally the resulting permuted covariance matrix was subject to SVD and the permuted crossblock covariance was compared to the observed, non-permuted value.

First, we repeated the PLS analysis after randomly dropping all but one subject from each family, leaving 1,722 subjects. This yielded a single statistically significant latent variable (*p* = .0017, covariance=48%) with stable loadings on the general and ADHD factors (Supplementary Figure 1a) and brain loadings significantly correlated with the un-adjusted model (*r_s_* = .98, Supplementary Figure 2).

With these sampling weights (*76*), PLS revealed a single statistically significant latent variable (*p* = .04, covariance=38%) with stable loadings on the general, ADHD, and internalizing factors (Supplementary Figure 1b) and brain loadings significantly correlated with the un-adjusted model (*r_s_* = .87, Supplementary Figure 2).

Finally, we sought to confirm that the observed Hurst-Psychopathology gradient is not specific to the bifactor model’s characterization of psychopathology. Thus, we repeated the PLS analysis with t-scores derived from the 11 CBCL syndrome scale scores (Anxiety/Depression, Withdrawn/Depression, Somatic, Social, Thought, Attention, Rule Breaking, Aggressive, Internal, External, Total Problems, (*77*)), 6 CBCL DSM5 scale scores (Depression, Anxiety/Disordered, Somatic, ADHD, Oppositional, Conduct, (*78*)), and 3 CBCL 2007 scale scores (Sluggish Cognitive Tempo, Obsessive-Compulsive, Stress) (*79*). PLS revealed two statistically significant latent variables, the first of which (*p* = .017, covariance=66%) had stable positive loadings on all derived CBCL scores (Supplementary Figure 1c) and brain loadings significantly correlated with the un-adjusted model (*r_s_* = .84, Supplementary Figure 2).

### Spatial Null Model

Spatial auto-correlation-preserving permutation tests were used to assess statistical significance of correlations between Neurosynth terms and the Hurst-Psychopathology gradient. These tests, termed “spin tests”, are necessary since standard permutation tests which assume no spatial auto-correlation significantly inflate false positive rates (*80*). We used the BrainSMASH python package (https://brainsmash.readthedocs.io/en/latest/) to generate parametric null brain maps with preserved spatial auto-correlation (SA) structure (*81*). In short, BrainSMASH produces SA-preserving random maps whose variograms approximately match the variogram of an input brain map. Variograms are functions of spatial distance, d, which quantify the variance between all pairs of points that are a distance d away from each other. Here we use Euclidean distance calculated between the centroids of brain parcels.

### Functional Activation

Functional activations in the Destrieux 148 parcel cortical atlas (*82*) were obtained from ABCD Release2.0.1 tabulated imaging data available on the NIMH data archive (https://nda.nih.gov/abcd). After preprocessing, beta weights for a linear contrast between 2-back and 0-back blocks were computed by generalized linear model with motion estimates, derivatives, squared estimates, and squared derivatives included as nuisance regressors (*33*). The hemodynamic response function was modeled as gamma functions with temporal derivatives and convolved with square waves indicating each block. Average beta coefficients across runs of the 2-back vs. 0-back contrast were used to assess functional activation. These data were retrieved from the *abcd_tfncr1bwdp201.txt* and *abcd_tfncr1bwdp201.txt* files.

To compare the correlation between activations and gradient-scores to the Hurst-Psychopathology gradient, the activations were resampled via averaging into the 392 parcels of the Craddock atlas with the nilearn (*83*) function NiftiLabelsMasker.

## Acknowledgments

Data used in the preparation of this article were obtained from the Adolescent Brain Cognitive DevelopmentSM (ABCD) Study (https://abcdstudy.org), held in the NIMH Data Archive (NDA). This is a multisite, longitudinal study designed to recruit more than 10,000 children age 9-10 and follow them over 10 years into early adulthood. The ABCD Study® is supported by the National Institutes of Health and additional federal partners under award numbers U01DA041048, U01DA050989, U01DA051016, U01DA041022, U01DA051018, U01DA051037, U01DA050987, U01DA041174, U01DA041106, U01DA041117, U01DA041028, U01DA041134, U01DA050988, U01DA051039, U01DA041156, U01DA041025, U01DA041120, U01DA051038, U01DA041148, U01DA041093, U01DA041089, U24DA041123, U24DA041147. A full list of supporters is available at https://abcdstudy.org/federal-partners.html. A listing of participating sites and a complete listing of the study investigators can be found at https://abcdstudy.org/consortium members/.

ABCD consortium investigators designed and implemented the study and/or provided data but did not necessarily participate in the analysis or writing of this report. This manuscript reflects the views of the authors and may not reflect the opinions or views of the NIH or ABCD consortium investigators.

The ABCD data repository grows and changes over time. The ABCD data used in this report came from DOI 10.15154/1503209 and DOI 10.15154/1519007. DOIs can be found at https://dx.doi.org/10.15154/1503209 and https://dx.doi.org/10.15154/1519007.

This research also benefited from the ABCD Workshops on Brain Development, Mental Health, and Longitudinal Modeling, supported by the NIMH and NIH under award numbers R25MH120869, R25MH125545, and UG3DA045251. This work was partially supported by SCC-1952050 (to M.G.B.), R00MH117274 (to A.N.K.), and UG3DA045251 (to B.B.L.).

## Code Availability

Analysis code is available at https://github.com/enlberman/fractalpsychopyabcd

## Footnotes

1 See see (*11*), however, for a discussion of systems with high *H* in the absence of criticality

2 Values of *H* below 0.5 are possible for time series with negative autocorrelation. Likewise values of *H* above 1 are possible for multi-fractal and non-stationary timeseries. However, these values are rarely observed in fMRI data.

## References

1. Mandelbrot, B. How long is the coast of britain? statistical self-similarity and fractional dimension. science 156, 636–638 (1967).

2. Suckling, J., Wink, A. M., Bernard, F. A., Barnes, A. & Bullmore, E. Endogenous multifractal brain dynamics are modulated by age, cholinergic blockade and cognitive performance. Journal of neuroscience methods 174, 292–300 (2008).

3. He, B. J. Scale-free properties of the functional magnetic resonance imaging signal during rest and task. Journal of Neuroscience 31, 13786–13795 (2011).

4. Churchill, N. W. et al. Scale-free brain dynamics under physical and psychological distress: Pre-treatment effects in women diagnosed with breast cancer. Human Brain Mapping 36, 1077–1092 (2015).

5. Kardan, O. et al. Distinguishing cognitive effort and working memory load using scale-invariance and alpha suppression in eeg. NeuroImage 211, 116622 (2020).

6. Peng, C.-K. et al. Mosaic organization of dna nucleotides. Physical review e 49, 1685 (1994).

7. Kardan, O. et al. Scale-invariance in brain activity predicts practice effects in cognitive performance. bioRxiv (2020).

8. Wei, M. et al. Identifying major depressive disorder using Hurst exponent of resting-state brain networks. Psychiatry Research: Neuroimaging 214, 306–312 (2013). URL https://linkinghub.elsevier.com/retrieve/pii/S0925492713002643.

9. Akhrif, A., Romanos, M., Domschke, K., Schmitt-Boehrer, A. & Neufang, S. Fractal analysis of bold time series in a network associated with waiting impulsivity. Frontiers in physiology 9, 1378 (2018).

10. Cocchi, L., Gollo, L. L., Zalesky, A. & Breakspear, M. Criticality in the brain: A synthesis of neurobiology, models and cognition. Progress in neurobiology 158, 132–152 (2017).

11. Touboul, J. & Destexhe, A. Power-law statistics and universal scaling in the absence of criticality. Physical Review E 95, 012413 (2017).

12. Roberts, J. A., Iyer, K. K., Vanhatalo, S. & Breakspear, M. Critical role for resource constraints in neural models. Frontiers in systems neuroscience 8, 154 (2014).

13. Kinouchi, O. & Copelli, M. Optimal dynamical range of excitable networks at criticality. Nature physics 2, 348–351 (2006).

14. Gautam, S. H., Hoang, T. T., McClanahan, K., Grady, S. K. & Shew, W. L. Maximizing sensory dynamic range by tuning the cortical state to criticality. PLoS computational biology 11, e1004576 (2015).

15. Boedecker, J., Obst, O., Lizier, J. T., Mayer, N. M. & Asada, M. Information processing in echo state networks at the edge of chaos. Theory in Biosciences 131, 205–213 (2012).

16. Shriki, O. et al. Neuronal avalanches in the resting meg of the human brain. Journal of Neuroscience 33, 7079–7090 (2013).

17. Shriki, O. & Yellin, D. Optimal information representation and criticality in an adaptive sensory recurrent neuronal network. PLoS computational biology 12, e1004698 (2016).

18. Medaglia, J. D., Pasqualetti, F., Hamilton, R. H., Thompson-Schill, S. L. & Bassett, D. S. Brain and cognitive reserve: Translation via network control theory. Neuroscience & Biobehavioral Reviews 75, 53–64 (2017). URL https://linkinghub.elsevier.com/retrieve/pii/S0149763416302329.

19. Levens, S. M., Muhtadie, L. & Gotlib, I. H. Rumination and impaired resource allocation in depression. Journal of Abnormal Psychology 118, 757–766 (2009). URL http://doi.apa.org/getdoi.cfm?doi=10.1037/a0017206.

20. Jones, N. P., Siegle, G. J., Muelly, E. R., Haggerty, A. & Ghinassi, F. Poor performance on cognitive tasks in depression: Doing too much or not enough? *Cognitive, Affective*, & Behavioral Neuroscience 10, 129–140 (2010). URL http://link.springer.com/10.3758/CABN.10.1.129.

21. Moore, T. M. et al. Criterion validity and relationships between alternative hierarchical dimensional models of general and specific psychopathology. Journal of abnormal psychology 129, 677 (2020).

22. Gao, W., Chen, S., Biswal, B., Lei, X. & Yuan, J. Temporal dynamics of spontaneous default-mode network activity mediate the association between reappraisal and depression. Social cognitive and affective neuroscience 13, 1235–1247 (2018).

23. Čukić, M. et al. Nonlinear analysis of eeg complexity in episode and remission phase of recurrent depression. International journal of methods in psychiatric research 29, e1816 (2020).

24. Ide, J. S., Hu, S., Zhang, S., Mujica-Parodi, L. R. & Chiang-shan, R. L. Power spectrum scale invariance as a neural marker of cocaine misuse and altered cognitive control. NeuroImage: Clinical 11, 349–356 (2016).

25. Sokunbi, M. O. Children with adhd exhibit lower fmri spectral exponent than their typically developing counterparts (Organisation for Human Brain Mapping (OHBM), USA., 2018).

26. Sokunbi, M. O. et al. Nonlinear complexity analysis of brain fmri signals in schizophrenia. Plos one 9, e95146 (2014).

27. Lai, M.-C. et al. A Shift to Randomness of Brain Oscillations in People with Autism. Biological Psychiatry 68, 1092–1099 (2010). URL https://linkinghub.elsevier.com/retrieve/pii/S0006322310007080.

28. Caspi, A. et al. The p factor: one general psychopathology factor in the structure of psychiatric disorders? Clinical psychological science 2, 119–137 (2014).

29. Lahey, B. B. et al. Is there a general factor of prevalent psychopathology during adulthood? Journal of abnormal psychology 121, 971 (2012).

30. Lahey, B. B., Krueger, R. F., Rathouz, P. J., Waldman, I. D. & Zald, D. H. A hierarchical causal taxonomy of psychopathology across the life span. Psychological bulletin 143, 142 (2017).

31. Kaczkurkin, A. N. et al. Approaches to defining common and dissociable neurobiological deficits associated with psychopathology in youth. Biological psychiatry 88, 51–62 (2020).

32. Reise, S. P. The rediscovery of bifactor measurement models. Multivariate behavioral research 47, 667–696 (2012).

33. Hagler Jr, D. J. et al. Image processing and analysis methods for the adolescent brain cognitive development study. Neuroimage 202, 116091 (2019).

34. Craddock, R. C., James, G. A., Holtzheimer III, P. E., Hu, X. P. & Mayberg, H. S. A whole brain fmri atlas generated via spatially constrained spectral clustering. Human brain mapping 33, 1914–1928 (2012).

35. Rosenberg, M. D. et al. Behavioral and neural signatures of working memory in childhood. Journal of Neuroscience (2020).

36. Harden, K. P. et al. Genetic Associations Between Executive Functions and a General Factor of Psychopathology. Journal of the American Academy of Child & Adolescent Psychiatry 59, 749–758 (2020). URL https://linkinghub.elsevier.com/retrieve/pii/S0890856719303272.

37. Huang-Pollock, C., Shapiro, Z., Galloway-Long, H. & Weigard, A. Is Poor Working Memory a Transdiagnostic Risk Factor for Psychopathology? Journal of Abnormal Child Psychology 45, 1477–1490 (2017). URL http://link.springer.com/10.1007/s10802-016-0219-8.

38. Reininghaus, U. et al. Reasoning bias, working memory performance and a transdiagnostic phenotype of affective disturbances and psychotic experiences in the general population. Psychological Medicine 49, 1799–1809 (2019).

39. Weissman, D. G. et al. Difficulties with emotion regulation as a transdiagnostic mechanism linking child maltreatment with the emergence of psychopathology. Development and Psychopathology 31, 899–915 (2019).

40. Keenan, K. Emotion Dysregulation as a Risk Factor for Child Psychopathology. Clinical Psychology: Science and Practice 7, 418–434 (2006). URL http://doi.wiley.com/10.1093/clipsy.7.4.418.

41. Wills, T. A., Simons, J. S., Sussman, S. & Knight, R. Emotional self-control and dysregulation: A dual-process analysis of pathways to externalizing/internalizing symptomatology and positive well-being in younger adolescents. Drug and Alcohol Dependence 163, S37–S45 (2016). URL https://linkinghub.elsevier.com/retrieve/pii/S0376871616001198.

42. McIntosh, A. R. & Lobaugh, N. J. Partial least squares analysis of neuroimaging data: applications and advances. Neuroimage 23, S250–S263 (2004).

43. Krishnan, A., Williams, L. J., McIntosh, A. R. & Abdi, H. Partial least squares (pls) methods for neuroimaging: a tutorial and review. Neuroimage 56, 455–475 (2011).

44. Heeringa, S. G. & Berglund, P. A. A guide for population-based analysis of the adolescent brain cognitive development (abcd) study baseline data. BioRxiv (2020).

45. Dick, A. S. et al. Meaningful associations in the adolescent brain cognitive development study. NeuroImage 118262 (2021).

46. Weintraub, S. et al. Cognition assessment using the nih toolbox. Neurology 80, S54–S64 (2013).

47. Yarkoni, T., Poldrack, R. A., Nichols, T. E., Van Essen, D. C. & Wager, T. D. Large-scale automated synthesis of human functional neuroimaging data. Nature methods 8, 665–670 (2011).

48. Hansen, J. Y. et al. Molecular signatures of cognition and affect. bioRxiv (2020).

49. Poldrack, R. A. et al. The cognitive atlas: toward a knowledge foundation for cognitive neuroscience. Frontiers in neuroinformatics 5, 17 (2011).

50. Shafiei, G. et al. Topographic gradients of intrinsic dynamics across neocortex. Elife 9, e62116 (2020).

51. Kardan, O. et al. Adult neuromarkers of sustained attention and working memory predict inter- and intra-individual differences in these processes in youth. bioRxiv (2021). URL https://www.biorxiv.org/content/early/2021/08/02/2021.08.01.454530. https://www.biorxiv.org/content/early/2021/08/02/2021.08.01.454530.full.

52. Cerquera, A., Arns, M., Buitrago, E., Gutiérrez, R. & Freund, J. Nonlinear dynamics measures applied to eeg recordings of patients with attention deficit/hyperactivity disorder: quantifying the effects of a neurofeedback treatment. In 2012 Annual International Conference of the IEEE Engineering in Medicine and Biology Society, 1057–1060 (IEEE, 2012).

53. Long, Z. et al. A Brainnetome Atlas Based Mild Cognitive Impairment Identification Using Hurst Exponent. Frontiers in Aging Neuroscience 10, 103 (2018). URL http://journal.frontiersin.org/article/10.3389/fnagi.2018.00103/full.

54. de Arcangelis, L., Perrone-Capano, C. & Herrmann, H. J. Self-Organized Criticality model for Brain Plasticity. Physical Review Letters 96, 028107 (2006). URL http://arxiv.org/abs/q-bio/0602014. ArXiv: q-bio/0602014.

55. Kitzbichler, M. G., Smith, M. L., Christensen, S. R. & Bullmore, E. Broadband criticality of human brain network synchronization. PLoS Comput Biol 5, e1000314 (2009).

56. Jolly, E. & Chang, L. J. The flatland fallacy: Moving beyond low–dimensional thinking. Topics in cognitive science 11, 433–454 (2019).

57. Bassett, D. S., Zurn, P. & Gold, J. I. On the nature and use of models in network neuroscience. Nature Reviews Neuroscience 19, 566–578 (2018).

58. Esteban, O. et al. fmriprep: a robust preprocessing pipeline for functional mri. Nature methods 16, 111–116 (2019).

59. Gorgolewski, K. et al. Nipype: a flexible, lightweight and extensible neuroimaging data processing framework in python. Frontiers in neuroinformatics 5, 13 (2011).

60. Gulban, O. F. et al. poldracklab/pydeface: v2.0.0 (2019). URL https://doi.org/10.5281/zenodo.3524401.

61. Tustison, N. J. et al. N4itk: improved n3 bias correction. IEEE transactions on medical imaging 29, 1310–1320 (2010).

62. Avants, B. B., Epstein, C. L., Grossman, M. & Gee, J. C. Symmetric diffeomorphic image registration with cross-correlation: evaluating automated labeling of elderly and neurodegenerative brain. Medical image analysis 12, 26–41 (2008).

63. Zhang, Y., Brady, M. & Smith, S. Segmentation of brain mr images through a hidden markov random field model and the expectation-maximization algorithm. IEEE transactions on medical imaging 20, 45–57 (2001).

64. Greve, D. N. & Fischl, B. Accurate and robust brain image alignment using boundary-based registration. Neuroimage 48, 63–72 (2009).

65. Power, J. D. et al. Methods to detect, characterize, and remove motion artifact in resting state fmri. Neuroimage 84, 320–341 (2014).

66. Power, J. D., Barnes, K. A., Snyder, A. Z., Schlaggar, B. L. & Petersen, S. E. Spurious but systematic correlations in functional connectivity mri networks arise from subject motion. Neuroimage 59, 2142–2154 (2012).

67. Power, J. D. et al. Methods to detect, characterize, and remove motion artifact in resting state fmri. Neuroimage 84, 320–341 (2014).

68. Cox, R. W. Afni: software for analysis and visualization of functional magnetic resonance neuroimages. Computers and Biomedical research 29, 162–173 (1996).

69. Churchill, N. W. et al. The suppression of scale-free fmri brain dynamics across three different sources of effort: aging, task novelty and task difficulty. Scientific reports 6, 30895 (2016).

70. Barch, D. M. et al. Demographic, physical and mental health assessments in the adolescent brain and cognitive development study: Rationale and description. Developmental cognitive neuroscience 32, 55–66 (2018).

71. Clark, D. A. et al. The general factor of psychopathology in the adolescent brain cognitive development (abcd) study: A comparison of alternative modeling approaches. Clinical Psychological Science 9, 169–182 (2021).

72. Class, Q. A. et al. Socioemotional dispositions of children and adolescents predict general and specific second-order factors of psychopathology in early adulthood: A 12-year prospective study. Journal of abnormal psychology 128, 574 (2019).

73. Hastie, T. & Qian, J. Glmnet vignette. Retrieved June 9, 1–30 (2014).

74. Hastie, T., Tibshirani, R. & Friedman, J. The elements of statistical learning. Cited on 33 (2009).

75. Becker, J.-M. Weighted partial least squares–a new method to account for sampling weights in pls path modeling.

76. Garavan, H. et al. Recruiting the abcd sample: Design considerations and procedures. Developmental cognitive neuroscience 32, 16–22 (2018).

77. Achenbach, T. M., Ruffle, T. M. et al. The child behavior checklist and related forms for assessing behavioral/emotional problems and competencies. Pediatrics in review 21, 265–271 (2000).

78. Achenbach, T. M., Dumenci, L. & Rescorla, L. A. Dsm-oriented and empirically based approaches to constructing scales from the same item pools. Journal of clinical child and adolescent psychology 32, 328–340 (2003).

79. Achenbach, T. M. & Rescorla, L. Achenbach system of empirically based assessment. Retrieved from Mental Measurements Yearbook via EBSCOhost (2007).

80. Markello, R. & Misic, B. Comparing spatially-constrained null models for parcellated brain maps. BioRxiv (2020).

81. Burt, J. B., Helmer, M., Shinn, M., Anticevic, A. & Murray, J. D. Generative modeling of brain maps with spatial autocorrelation. NeuroImage 220, 117038 (2020).

82. Destrieux, C., Fischl, B., Dale, A. & Halgren, E. Automatic parcellation of human cortical gyri and sulci using standard anatomical nomenclature. Neuroimage 53, 1–15 (2010).

83. Abraham, A. et al. Machine learning for neuroimaging with scikit-learn. Frontiers in neuroinformatics 8, 14 (2014).

