## Supplementary Figures and Tables for "A Scale-Free Gradient of Cognitive Resource Disruptions in Childhood Psychopathology"

### Supplementary materials

#### Supplementary Figures

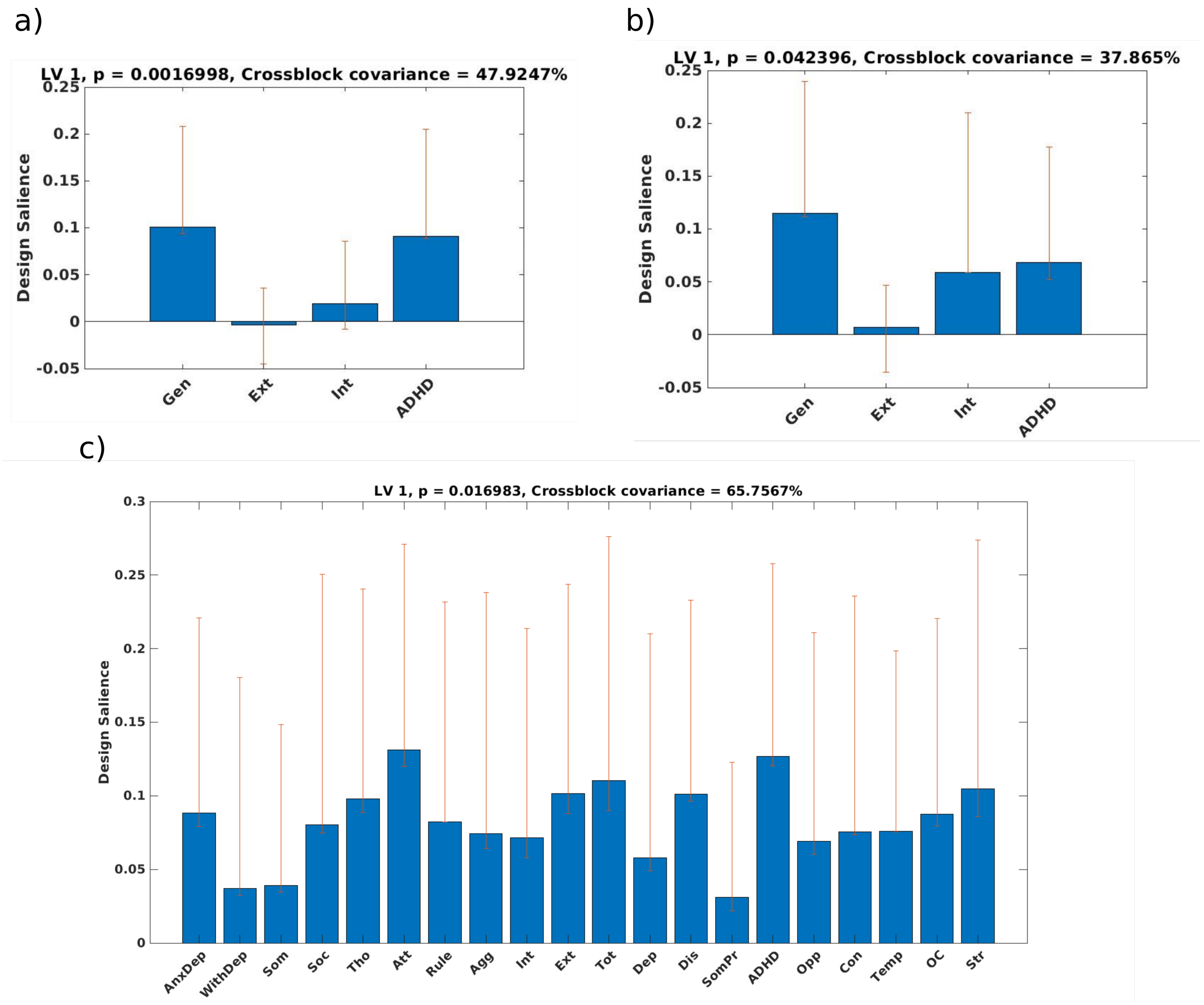


Supplementary Figure 1: Design loadings for PLS sensitivity models. These plots show the behavioral loadings on the significant latent variable for each of the sensitivity tests. a) Only one member of each family was retained. b) Non-participation and post-stratification weights were applied. c) CBCL scales were used instead of extracted bifactor scores. AnxDep = Anxiety Depression Syndrome Scale; WithDep = Withdrawn Depression Syndrome Scale; Som = Somatic Syndrome Scale; Soc = Social Syndrome Scale; Tho = Thought Syndrome Scale; Att = Attention Syndrome Scale; Rule = Rule Breaking Syndrome Scale; Agg = Aggressive Syndrome Scale; Int = Internal Syndrome Scale; Ext = External Syndrome Scale; Tot = Total Problems Syndrome Scale; Dep = Depression DSM5 Scale; Dis = Anxiety/Disordered DSM5 Scale; SomPr = Somatic Problems DSM5 Scale; ADHD = ADHD DSM5 Scale; Opp = Oppositional DSM5 Scale; Con = Conduct DSM5 Scale; Temp = Sluggish Cognitive Tempo 2007 Scale; OC = Obsessive-Compulsive 2007 Scale; Str = Stress 2007 Scale.

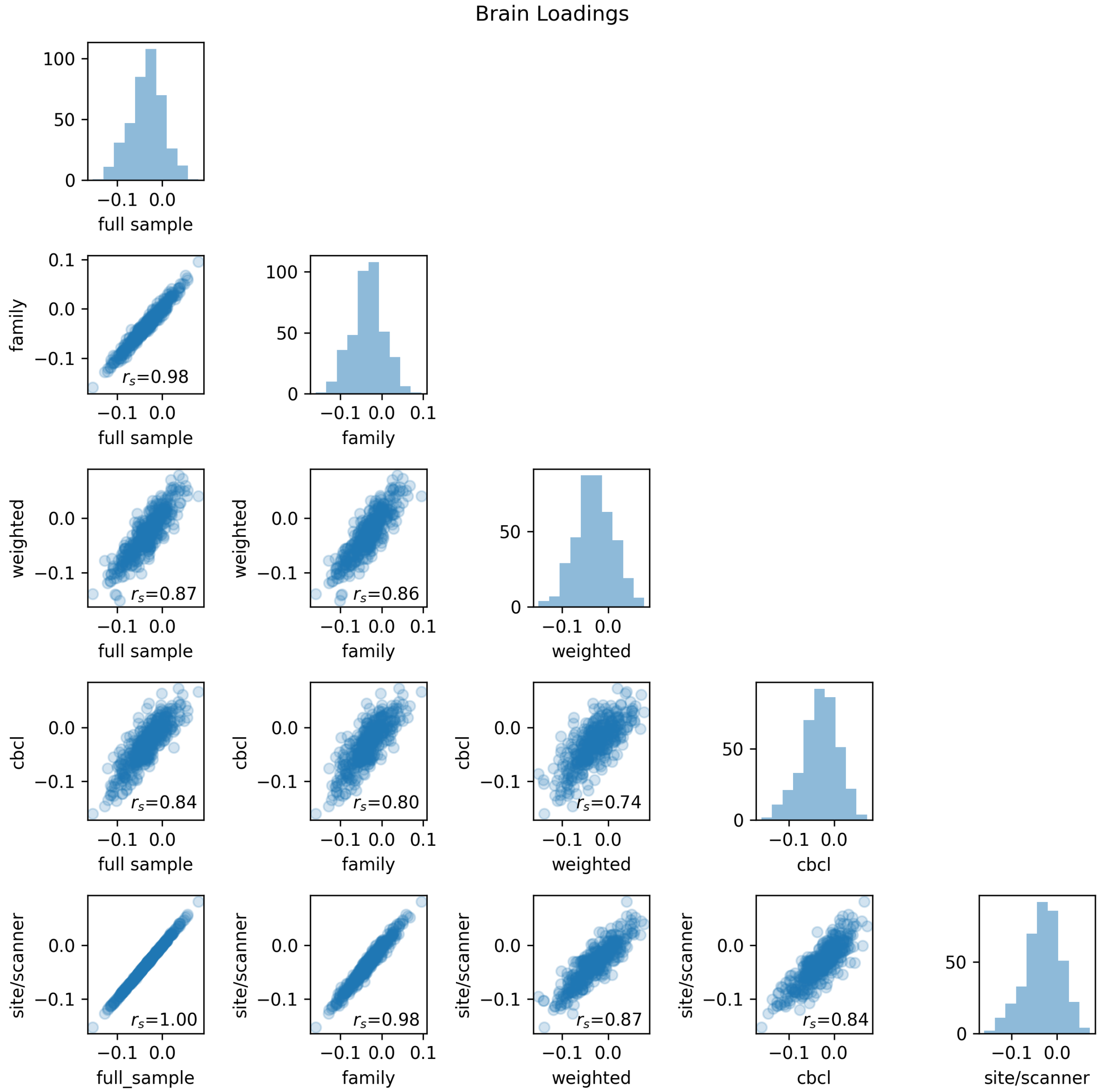


Supplementary Figure 2: Comparison of brain loadings between the un-adjusted full sample model, the model where only one family member was retained, the model adjusted for post-stratification and non-participation, the model with CBCL scores instead of extracted bifactor scores, and the model where data was residualized with respect to site and scanner. All correlations are significant.

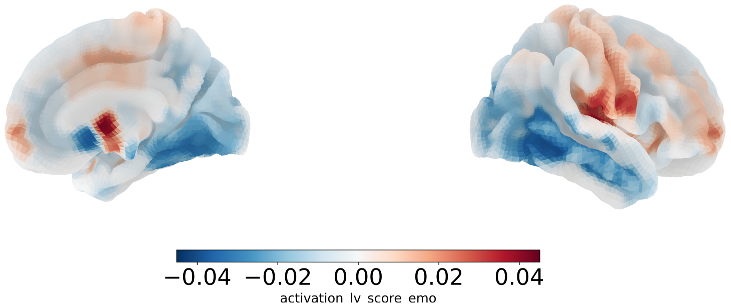


Supplementary Figure 3: Correlations between the Hurst-Psychopathology gradient and functional activations defined from a contrast between emotional and neutral faces. Functional activations were not significantly correlated to the H-P gradient after multiple comparison correction.

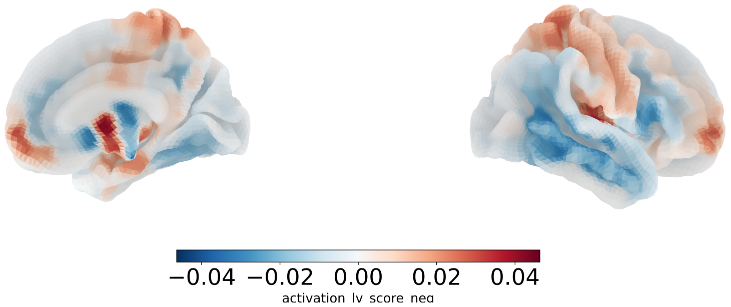


Supplementary Figure 4: Correlations between the Hurst-Psychopathology gradient and functional activations defined from a contrast between negative and neutral faces. Functional activations were not significantly correlated to the H-P gradient after multiple comparison correction.

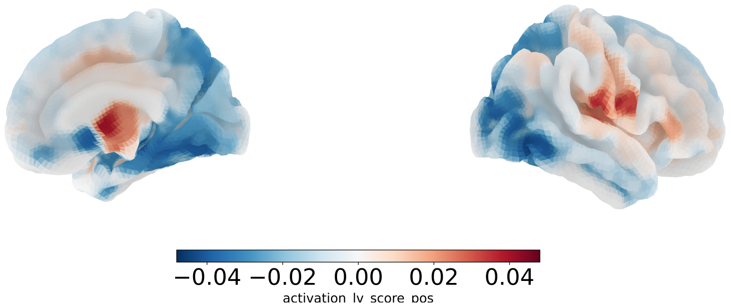


Supplementary Figure 5: Correlations between the Hurst-Psychopathology gradient and functional activations defined from a contrast between positive and neutral faces. Functional activations were not significantly correlated to the H-P gradient after multiple comparison correction.

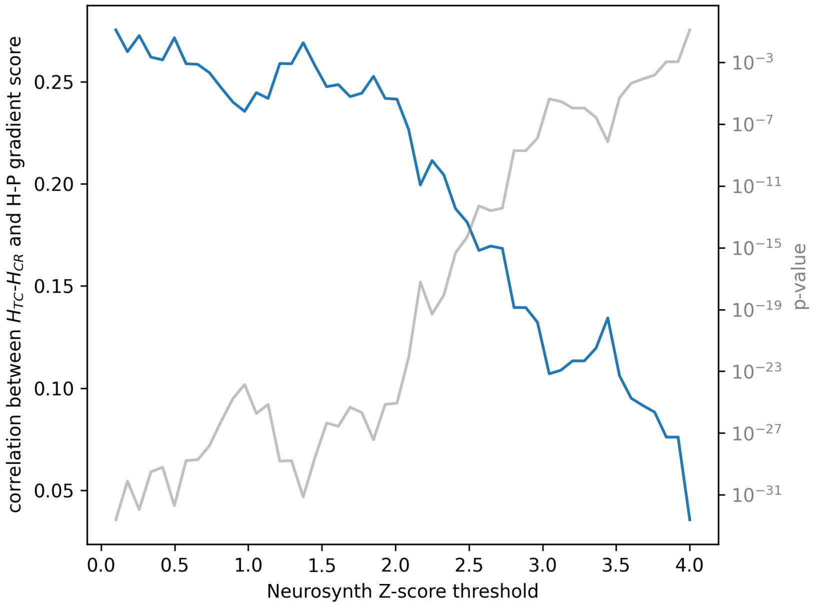


Supplementary Figure 6: Psychopathology is associated with relatively higher Hurst exponents in Task-Cognition areas relative to Cue-Response areas. The difference between the mean Hurst exponent for Task-Cognition ($H_{TC}$) and Cue-Response ($H_{CR}$) was correlated with subjects' gradient scores (i.e., the degree to which each subject exemplifies the H-P gradient during the EN-back task). This resulted in a significant positive correlation across all choices of Z-score threshold.

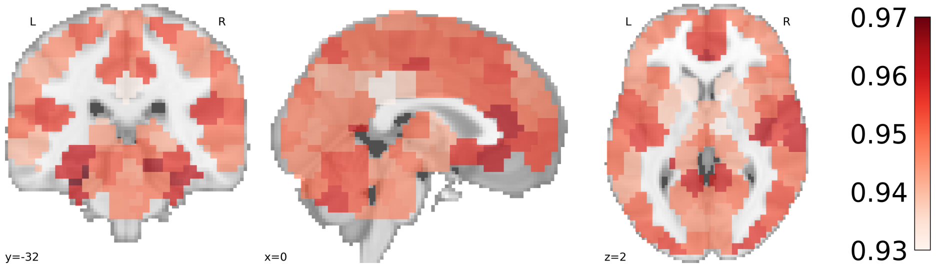


Supplementary Figure 7: Hurst $R^2$, mean for all subjects. High $R^2$ values on average demonstrate good fits in the detrended fluctuation analysis, and suggest that the fMRI time series in these data are well described by a mono-fractal process, as opposed to a multi-fractal process.

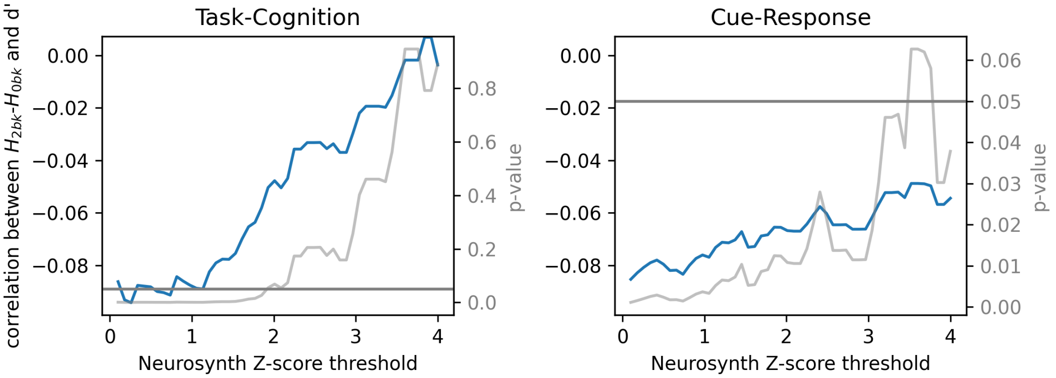


Supplementary Figure 8: Correlations between block level 2- vs. 0-back $H$ contrast and d'. Correlations (blue) were computed across all choices of Z-score threshold used to define the task-cognition and cue-response ROIs from Neurosynth meta-analysis probabilistic activation maps. Significance of the correlations were assessed at all choices of threshold (gray). The p=0.05 level is show as the gray horizontal line. We found evidence associating suppression of $H$ with d' in both task-cognition and cue-response areas.

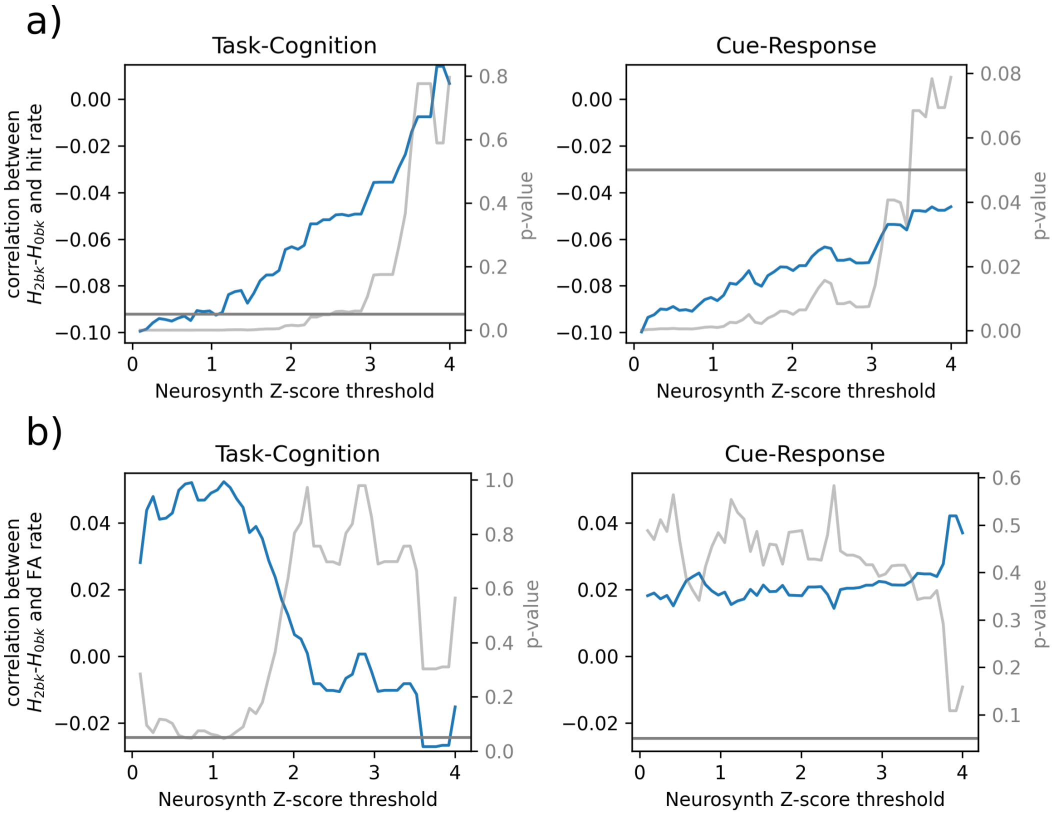


Supplementary Figure 9: Correlations between block level 2- vs. 0-back $H$ contrast and hit rate (a, top) and false alarm rate, FA (b, bottom). Correlations (blue) were computed across all choices of Z-score threshold used to define the task-cognition and cue-response ROIs from Neurosynth meta-analysis probabilistic activation maps. Significance of the correlations were assessed at all choices of threshold (gray). The p=0.05 level is show as the gray horizontal line. a/top) We found evidence associating suppression of $H$ with hit rate in both task-cognition and cue-response areas. b/bottom) We found no evidence of a relationship between the 2- vs. 0-back $H$ contrast and FA rate in neither task-cognition nor cue-response areas.

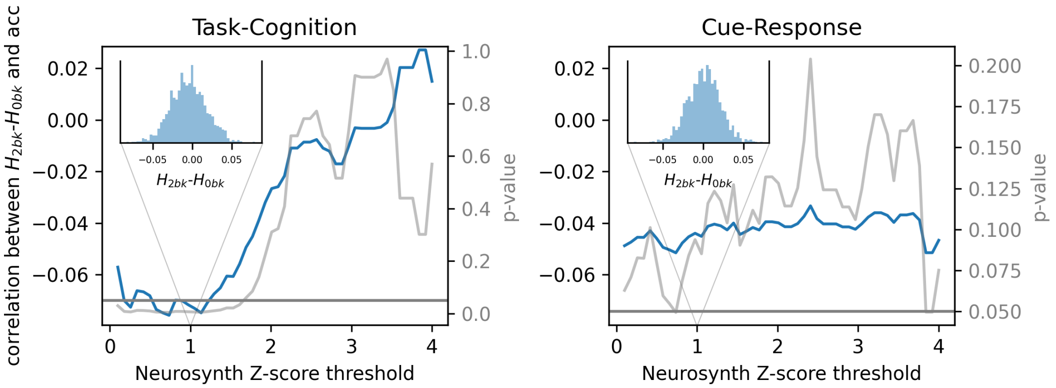


Supplementary Figure 10: Correlations between block level 2- vs. 0-back $H$ contrast and 2-back accuracy (acc). Correlations (blue) were computed across all choices of Z-score threshold used to define the task-cognition and cue-response ROIs from Neurosynth meta-analysis probabilistic activation maps. Significance of the correlations were assessed at all choices of threshold (gray). The p=0.05 level is show as the gray horizontal line. We found evidence associating suppression of $H$ with acc in only task-cognition areas. Insets show the distribution of the 2-back vs. 0-back contrast for a Z-score threshold of 1.

#### Supplementary Tables

| | Included Participants | | | Excluded Participants | | | Differences | | |
|---|---|---|---|---|---|---|---|---|---|
| | Mean | SE | N | Mean | SE | N | t | d.f. | p < |
| Age in months | 120.3 | 0.18 | 1839 | 118.7 | 0.07 | 10039 | **8.5** | | **2.075e-17** |
| Household size | 4.9 | 0.05 | 1818 | 4.7 | 0.02 | 9781 | **3.77** | | **0.0002** |
| | n | % | N | n | % | N | $\chi^2$ | d.f. | p < |
| Sex (% female) | 1018 | 0.55 | 1839 | 4664 | 0.46 | 10039 | **48.95** | **1** | **2.6e-12** |
| Race-ethnicity | | | 1839 | | | 10037 | **146.74** | **4** | **1e-30** |
| Non - Hispanic white | 1176 | 0.64 | | 5006 | 0.5 | | | | |
| African - American | 148 | 0.08 | | 1636 | 0.16 | | | | |
| Hispanic | 311 | 0.17 | | 2100 | 0.21 | | | | |
| Asian | 39 | 0.02 | | 213 | 0.02 | | | | |
| Other | 165 | 0.09 | | 1082 | 0.11 | | | | |
| Income | | | 1839 | | | 10039 | **105.04** | **3** | **1.2e-22** |
| <50K | 347 | 0.19 | | 2877 | 0.29 | | | | |
| >50K & <100k | 834 | 0.45 | | 3731 | 0.37 | | | | |
| >100k | 541 | 0.29 | | 2530 | 0.25 | | | | |
| Not reported | 117 | 0.06 | | 901 | 0.09 | | | | |
| Maternal Education | | | 1839 | | | 10039 | **97.81** | **4** | **2.9e-20** |
| <HS Diploma | 58 | 0.03 | | 728 | 0.07 | | | | |
| HS Diploma/GED | 885 | 0.48 | | 3986 | 0.4 | | | | |
| Some College | 139 | 0.08 | | 1121 | 0.11 | | | | |
| College | 503 | 0.27 | | 2508 | 0.25 | | | | |
| Advanced Degree | 139 | 0.08 | | 1121 | 0.11 | | | | |
| Not reported | 503 | 0.27 | | 2508 | 0.25 | | | | |

Supplementary Table 1: Demographics of included and excluded participants. Tests in bold are significant after false discovery rate correction (adopting a 5% false discovery rate) for six tests.

| | | | |
|---|---|---|---|
| **Dep. Variable:** | List Sorting Performance | **R-squared:** | 0.101 |
| **Model:** | OLS | **Adj. R-squared:** | 0.100 |
| **Method:** | Least Squares | **F-statistic:** | 99.37 |
| **Date:** | Sun, 22 Aug 2021 | **Prob (F-statistic):** | 1.27e-41 |
| **Time:** | 22:08:54 | **Log-Likelihood:** | -6620.2 |
| **No. Observations:** | 1769 | **AIC:** | 1.325e+04 |
| **Df Residuals:** | 1766 | **BIC:** | 1.326e+04 |
| **Df Model:** | 2 | | |

| | **coef** | **std err** | **t** | **P> \|t\|** | **[0.025** | **0.975]** |
|---|---|---|---|---|---|---|
| **const** | 79.5768 | 3.200 | 24.866 | 0.000 | 73.300 | 85.854 |
| $H$ **in H-P gradient** | 19.6572 | 4.317 | 4.553 | 0.000 | 11.190 | 28.125 |
| **d' (in scanner)** | 3.1741 | 0.260 | 12.227 | 0.000 | 2.665 | 3.683 |

| | | | |
|---|---|---|---|
| **Omnibus:** | 65.434 | **Durbin-Watson:** | 2.015 |
| **Prob(Omnibus):** | 0.000 | **Jarque-Bera (JB):** | 73.182 |
| **Skew:** | -0.458 | **Prob(JB):** | 1.28e-16 |
| **Kurtosis:** | 3.392 | **Cond. No.** | 48.3 |

Supplementary Table 2: Linear model predicting out-of-scanner performance from in-scanner performance and $H$ in the H-P gradient network.

| | Included Participants | | | Excluded Participants | | | Differences | | |
|---|---|---|---|---|---|---|---|---|---|
| | Mean | SE | N | Mean | SE | N | t | | p < |
| Age in months | 120.2 | 0.25 | 888 | 120.5 | 0.25 | 951 | -0.76 | | 0.4459 |
| Household size | 4.9 | 0.1 | 880 | 4.8 | 0.05 | 938 | 1.33 | | 0.1850 |
| general factor | -0.0 | 0.03 | 888 | -0.1 | 0.03 | 951 | 1.35 | | 0.1773 |
| Internalizing factor | 0.1 | 0.02 | 888 | 0.1 | 0.02 | 951 | 0.9 | | 0.3697 |
| Externalizing factor | -0.1 | 0.02 | 888 | -0.1 | 0.02 | 951 | 0.05 | | 0.9585 |
| ADHD factor | -0.1 | 0.02 | 888 | -0.1 | 0.02 | 951 | 1.69 | | 0.0917 |
| | n | % | N | n | % | N | $\chi^2$ | d.f. | p < |
| Sex (% female) | 457 | 0.51 | 888 | 561 | 0.59 | 951 | **10.22** | **1** | **0.0014** |
| Race-ethnicity | | | 888 | | | 951 | 11.39 | 4 | 0.0225 |
| Non - Hispanic white | 592 | 0.67 | | 584 | 0.61 | | | | |
| African - American | 61 | 0.07 | | 87 | 0.09 | | | | |
| Hispanic | 150 | 0.17 | | 161 | 0.17 | | | | |
| Asian | 21 | 0.02 | | 18 | 0.02 | | | | |
| Other | 64 | 0.07 | | 101 | 0.11 | | | | |
| Income | | | 888 | | | 951 | 9.54 | 3 | 0.0229 |
| <50K | 179 | 0.2 | | 168 | 0.18 | | | | |
| >50K & <100k | 371 | 0.42 | | 463 | 0.49 | | | | |
| >100k | 282 | 0.32 | | 259 | 0.27 | | | | |
| Not reported | 56 | 0.06 | | 61 | 0.06 | | | | |
| Maternal Education | | | 888 | | | 951 | 10.91 | 4 | 0.0276 |
| <HS Diploma | 24 | 0.03 | | 34 | 0.04 | | | | |
| Hs Diploma/GED | 438 | 0.49 | | 447 | 0.47 | | | | |
| Some College | 67 | 0.08 | | 72 | 0.08 | | | | |
| College | 219 | 0.25 | | 284 | 0.3 | | | | |
| Advanced Degree | 67 | 0.08 | | 72 | 0.08 | | | | |
| Not reported | 219 | 0.25 | | 284 | 0.3 | | | | |

Supplementary Table 3: Demographics of study participants who had available 2-year follow up behavioral data for the in-scanner EN-back task. Tests in bold are significant after false discovery rate correction (adopting a 5% false discovery rate) for ten tests.

| | | | |
|---|---|---|---|
| **Dep. Variable:** | future d' | **R-squared:** | 0.243 |
| **Model:** | OLS | **Adj. R-squared:** | 0.240 |
| **Method:** | Least Squares | **F-statistic:** | 90.37 |
| **Date:** | Sun, 22 Aug 2021 | **Prob (F-statistic):** | 1.01e-50 |
| **Time:** | 22:08:31 | **Log-Likelihood:** | -1163.5 |
| **No. Observations:** | 849 | **AIC:** | 2335. |
| **Df Residuals:** | 845 | **BIC:** | 2354. |
| **Df Model:** | 3 | | |

| | **coef** | **std err** | **t** | **P> \|t\|** | **[0.025** | **0.975]** |
|---|---|---|---|---|---|---|
| **const** | -1.6641 | 0.502 | -3.317 | 0.001 | -2.649 | -0.679 |
| $H$ **in H-P gradient (baseline)** | 1.7510 | 0.584 | 2.998 | 0.003 | 0.605 | 2.897 |
| **d' (baseline)** | 0.4162 | 0.036 | 11.537 | 0.000 | 0.345 | 0.487 |
| **List sorting performance (baseline)** | 0.0207 | 0.003 | 6.258 | 0.000 | 0.014 | 0.027 |

| | | | |
|---|---|---|---|
| **Omnibus:** | 284.338 | **Durbin-Watson:** | 1.984 |
| **Prob(Omnibus):** | 0.000 | **Jarque-Bera (JB):** | 3034.181 |
| **Skew:** | -1.201 | **Prob(JB):** | 0.00 |
| **Kurtosis:** | 11.944 | **Cond. No.** | 2.23e+03 |

Supplementary Table 4: Linear model predicting future in-scanner performance from baseline in-scanner performance, out-of-scanner performance and $H$ in the H-P gradient network.

| lhs | op | rhs | est.std | se | z | pvalue | ci.lower | ci.upper |
|---|---|---|---|---|---|---|---|---|
| acc | ~ | bf1 | −.056 | .024 | −2.323 | .020 | −.103 | −.009 |
| acc | ~ | bf2 | .041 | .024 | 1.696 | .090 | −.006 | .089 |
| acc | ~ | bf3 | −.019 | .024 | −.774 | .439 | −.066 | .029 |
| acc | ~ | bf4 | −.051 | .023 | −2.181 | .029 | −.097 | −.005 |
| acc | ~ | h | .223 | .023 | 9.726 | 0 | .178 | .268 |
| h | ~ | bf1 | −.108 | .024 | −4.458 | .00001 | −.155 | −.060 |
| h | ~ | bf2 | −.010 | .025 | −.386 | .699 | −.058 | .039 |
| h | ~ | bf3 | −.026 | .025 | −1.060 | .289 | −.075 | .022 |
| h | ~ | bf4 | −.094 | .024 | −3.946 | .0001 | −.140 | −.047 |

Supplementary Table 5: Mediation with Hurst as the mediator. h is the Hurst exponent in the H-P gradient network. acc is 2-back accuracy. bf1-4 are extracted factor scores for, respectively, the general factor of psychopathology, the externalizing factor, the internalizing factor, and the ADHD factor.

| lhs | op | rhs | est.std | se | z | pvalue | ci.lower | ci.upper |
|---|---|---|---|---|---|---|---|---|
| acc | ~ | bf1 | −.056 | .023 | −2.385 | .017 | −.101 | −.010 |
| acc | ~ | bf2 | .041 | .023 | 1.782 | .075 | −.004 | .087 |
| acc | ~ | bf3 | −.019 | .023 | −.809 | .419 | −.064 | .027 |
| acc | ~ | bf4 | −.051 | .023 | −2.198 | .028 | −.097 | −.006 |
| acc | ~ | h | .223 | .023 | 9.842 | 0 | .178 | .267 |
| bf1 | ~ | h | −.114 | .024 | −4.839 | 0.00000 | −.160 | −.068 |
| bf2 | ~ | h | −.010 | .024 | −.402 | .688 | −.057 | .037 |
| bf3 | ~ | h | −.036 | .024 | −1.499 | .134 | −.083 | .011 |
| bf4 | ~ | h | −.093 | .024 | −3.924 | .0001 | −.140 | −.047 |

Supplementary Table 6: Mediation with extracted bifactor scores as the mediator.h is the Hurst exponent in the H-P gradient network. acc is 2-back accuracy. bf1-4 are extracted factor scores for, respectively, the general factor of psychopathology, the externalizing factor, the internalizing factor, and the ADHD factor.

Supplementary Table 7: Harvard-Oxford labels, PLS loadings, bootstrap ratios and MNI coordinates for brain regions with absolute value bootstrap ratios $> 2.5$.

| Harvard-Oxford Label | PLS loading | bootstrap ratio | MNI coordinates |
|---|---|---|---|
| Central Opercular Cortex | -0.1539 | -4.65 | (-58.0, -24.5, 18.5) |
| Superior Frontal Gyrus | -0.1278 | -4.0 | (24.7, 10.2, 57.5) |
| Inferior Frontal Gyrus, pars opercularis | -0.1209 | -3.71 | (-40.2, 8.1, 30.7) |
| Superior Frontal Gyrus | -0.1204 | -3.93 | (-21.9, 6.2, 63.3) |
| Middle Frontal Gyrus | -0.1159 | -3.57 | (28.5, -4.0, 52.8) |
| Precuneous Cortex | -0.1147 | -3.54 | (8.1, -66.5, 53.9) |
| Superior Frontal Gyrus | -0.1137 | -3.52 | (-16.3, -3.6, 68.0) |
| Superior Parietal Lobule | -0.1126 | -3.47 | (-16.7, -51.7, 66.0) |
| Middle Frontal Gyrus | -0.1114 | -3.53 | (-42.6, 22.5, 36.0) |
| Precentral Gyrus | -0.1091 | -3.41 | (-46.4, -0.5, 43.5) |
| Frontal Orbital Cortex | -0.1083 | -3.31 | (-27.2, 33.7, -15.9) |
| Superior Frontal Gyrus | -0.1083 | -3.33 | (20.7, -3.2, 66.3) |
| Central Opercular Cortex | -0.107 | -3.22 | (-41.6, -2.4, 8.1) |
| Cingulate Gyrus, anterior division | -0.1059 | -3.24 | (-7.8, -17.6, 43.7) |
| Postcentral Gyrus | -0.1042 | -3.33 | (62.9, -15.6, 17.9) |
| Postcentral Gyrus | -0.102 | -3.14 | (-27.0, -39.2, 63.9) |
| Central Opercular Cortex | -0.1017 | -3.08 | (41.6, -9.1, 13.9) |
| Cingulate Gyrus, posterior division | -0.0997 | -3.14 | (10.5, -27.3, 42.9) |

| | | | |
|---|---|---|---|
| Insular Cortex | -0.098 | -3.07 | (-35.7, -17.2, 1.1) |
| Lateral Occipital Cortex, superior division | -0.0968 | -2.93 | (-10.3, -65.8, 56.0) |
| Background | -0.0955 | -3.03 | (14.0, -56.1, -21.9) |
| Temporal Occipital Fusiform Cortex | -0.0948 | -3.02 | (37.3, -55.5, -17.2) |
| Frontal Pole | -0.0944 | -2.98 | (25.9, 59.6, -4.6) |
| Left Thalamus | -0.0942 | -2.94 | (-13.1, -23.1, 11.4) |
| Precentral Gyrus | -0.0923 | -2.8 | (-28.1, -6.4, 52.7) |
| Paracingulate Gyrus | -0.0913 | -2.76 | (-6.3, -0.7, 44.9) |
| Left Caudate | -0.0911 | -2.77 | (-13.2, 2.6, 16.3) |
| Lingual Gyrus | -0.0903 | -2.84 | (-24.3, -53.8, -9.2) |
| Precuneous Cortex | -0.0903 | -2.87 | (6.5, -52.8, 49.3) |
| Superior Parietal Lobule | -0.0894 | -2.82 | (26.5, -45.6, 64.5) |
| Postcentral Gyrus | -0.0893 | -2.75 | (-42.4, -18.4, 44.2) |
| Middle Frontal Gyrus | -0.0892 | -2.85 | (37.7, 3.0, 57.0) |
| Occipital Pole | -0.0888 | -2.75 | (-25.4, -95.0, 6.1) |
| Right Putamen | -0.0885 | -2.7 | (23.3, 9.0, 0.6) |
| Insular Cortex | -0.0885 | -2.79 | (33.1, 16.3, -8.0) |
| Cingulate Gyrus, posterior division | -0.0885 | -2.77 | (-5.4, -50.2, 19.6) |
| Background | -0.0872 | -2.76 | (18.6, -68.9, -27.2) |
| Frontal Pole | -0.0869 | -2.72 | (-45.8, 39.9, 4.1) |
| Occipital Pole | -0.0868 | -2.74 | (-8.5, -95.1, 0.5) |
| Left Putamen | -0.0859 | -2.54 | (-23.2, 7.9, 0.4) |
| Brain-Stem | -0.0847 | -2.6 | (-4.7, -25.1, -21.5) |
| Lateral Occipital Cortex, superior division | -0.0837 | -2.69 | (-27.3, -59.5, 56.7) |
| Background | -0.0832 | -2.58 | (29.9, -56.4, -27.9) |
| Superior Parietal Lobule | -0.0831 | -2.63 | (38.7, -46.0, 56.7) |
| Superior Parietal Lobule | -0.0829 | -2.64 | (40.5, -33.9, 44.7) |
| Juxtapositional Lobule Cortex | -0.0817 | -2.6 | (7.8, -13.6, 45.3) |
| Supramarginal Gyrus, anterior division | -0.0814 | -2.65 | (61.4, -22.8, 29.0) |
| Middle Frontal Gyrus | -0.0806 | -2.56 | (-36.2, 4.1, 56.0) |
| Frontal Pole | -0.0793 | -2.56 | (-25.8, 48.3, -14.0) |
